# Graphene Biointerface for Cardiac Arrhythmia Diagnosis and Treatment

**DOI:** 10.1101/2022.06.28.497825

**Authors:** Zexu Lin, Dmitry Kireev, Ning Liu, Shubham Gupta, Jessica LaPaino, Sofian N. Obaid, Zhiyuan Chen, Deji Akinwande, Igor R. Efimov

## Abstract

Heart rhythm disorders, known as arrhythmias, cause significant morbidity and are one of the leading causes of mortality. Cardiac arrhythmias are primarily treated by implantable devices, such as pacemakers and defibrillators, or by ablation therapy guided by electroanatomical mapping. Pharmacological treatments are mostly ineffective. Both implantable and ablation therapies require sophisticated biointerfaces for electrophysiological measurements of electrograms and delivery of therapeutic stimulation or ablation energy. In this work, we report for the first time on graphene biointerface for *in vivo* cardiac electrophysiology. Leveraging sub-micrometer thick tissue-conformable graphene arrays, we demonstrate sensing and stimulation of the open mammalian heart both *in vitro* and *in vivo.* Furthermore, we demonstrate graphene pacemaker treatment of a pharmacologically-induced arrhythmia, AV block. The arrays show effective electrochemical properties, namely interface impedance down to 40 Ohm×cm^2^ at 1kHz, charge storage capacity up to 63.7 mC/cm^2^, and charge injection capacity up to 704 μC/cm^2^. Transparency of the graphene structures allows for simultaneous optical mapping of cardiac action potentials and optogenetic stimulation while performing electrical measurements and stimulation. Our report presents evidence of the significant potential of graphene biointerfaces for the future clinical device- and catheter-based cardiac arrhythmias therapies.

## Introduction

Normal cardiac function is orchestrated by periodic electrical waves, commonly documented by an electrocardiogram (ECG)^1^, propagating from the natural pacemaker sinoatrial node to every electrically coupled myocyte^2^. Heart rhythm diseases (i.e., arrhythmia) could be deadly without instant electrical stimulation applied to the heart muscle^3^. Mechanical rigidity of lifesaving devices (Fig. S1), such as pacemakers and implantable cardioverter defibrillators, leads to non-ideal interfaces with soft tissues causing various complications (Table. S1)^4,5^. As a result, conformal devices melding to the curvilinear topology of the beating heart are desired.

Meanwhile, cardiac optophysiology and optogenetics are blossoming in basic scientific research both *ex vivo* and *in vivo* and have the potential to become powerful clinical tools^6–8^. Optical mapping offers unparalleled spatial resolution for probing physiological function (e.g., transmembrane potential, intracellular calcium, metabolism) with synthetic or endogenous (e.g., NADH and FAD) fluorescent reporters^9–11^. Optogenetics features unprecedented cell type-specific manipulation (e.g., cardiac pacing, arrhythmia termination) with a toolkit of opsins working across wide band light spectrum^6,12^. Conformal and transparent biointerfaces combining the advantages of electro- and opto-techniques and achieving simultaneous multimodal sensing and actuating would consequently benefit both basic scientific research and clinical therapy in studying and modulating complex cardiac events.

Emerging strategies in soft biointerfaces^13^ leverage materials and fabrication capabilities such as metal nanowires^14,15^, conductive fillers^16,17^, and serpentine architectures^18,19^. Issues of existing devices such as susceptibilities to oxidation and corrosion^20,21^, performance degradation in biological environments^22,23^, low conductivity^24,25^, high production cost^26,27^, non-transparency, and light-induced artifacts of metal electrodes^28^ leave room for improvement that utilizing intrinsically transparent materials with superior and stable electrical conductivity, biocompatibility, and mechanical property would be ideal. Table. 1 summarizes the state-of-the-art graphically, while the details can be found in Supplementary Tables. S1-4.

**Table 1.**
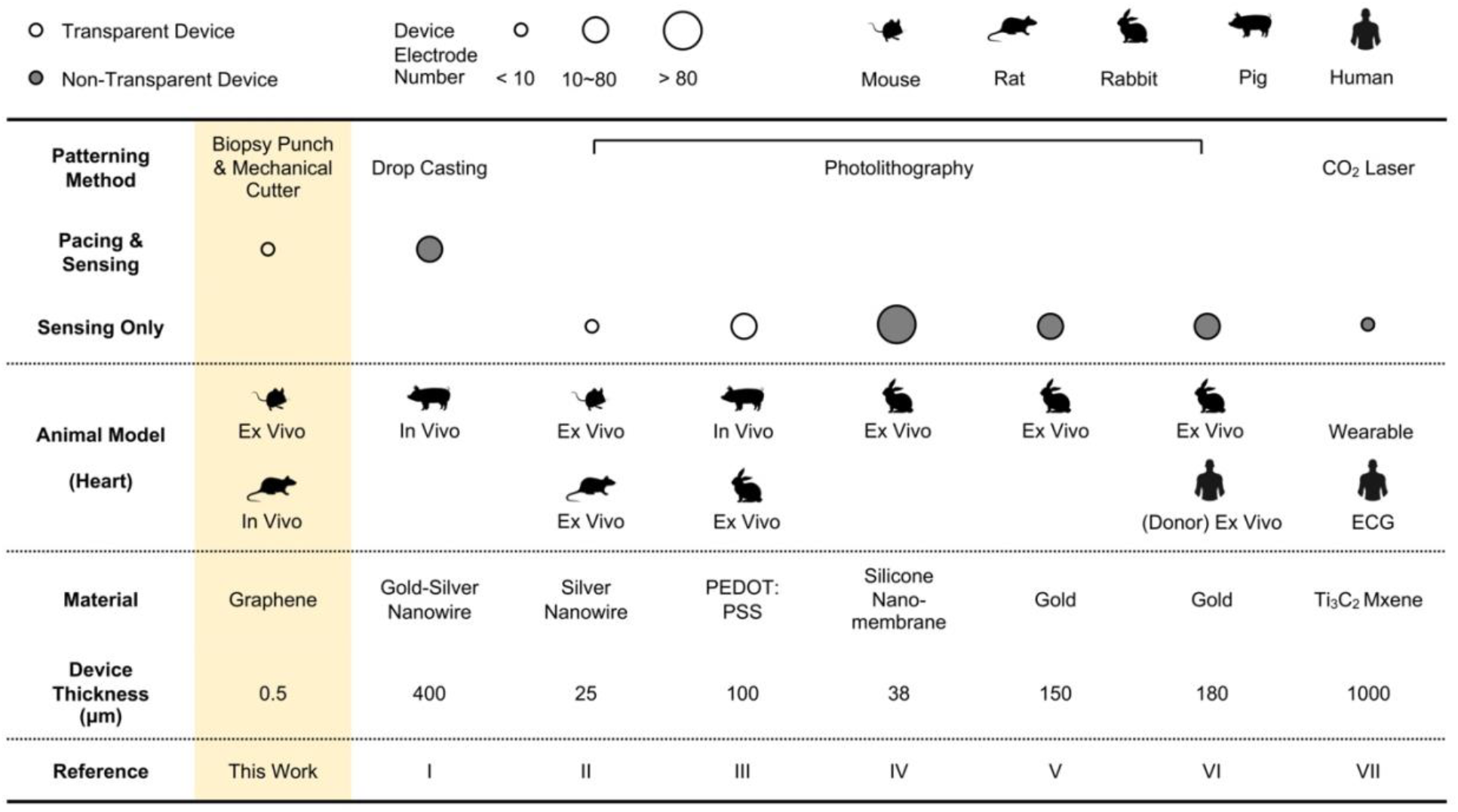
Examples of emergent soft biointerfaces for advanced cardiac electrophysiology. *Reference I*^15^, *II*^14^, *III*^17^, *IV*^41^, *V*^19^, *VI*^18^, *VII*^16^

Graphene possesses outstanding bioelectronics-relevant properties such as sub-nanometer thickness^29^, optical transparency (~97% for visible light)^30^, strength (130 GPa)^31^, stretchability (~15%)^32^, electrical conductivity^33^, and biocompatibility^34^. It has been reported to fabricate cardiopatches^35^, flexible microelectrodes^36^, and microprobes^37^ for electrical activities monitoring and stimulation in cardiac and neural cellular research (Table. S3). Studies also reported artifact-free optical recording and stimulation^28,38^ as well as MRI imaging compatibility^39^ through transparent graphene electrodes. However, all previous publications either focused on neuroscience or cultured cells and cardiac tissue that do not recapitulate in vivo cardiac biology. Recently, a cost- and time-effective fabrication protocol using off-the-shelf methods to create skin graphene electronic tattoos (GETs) of various shapes have been reported^40^. Combining these approaches could potentially help develop graphene-based biointerfaces for the intact heart, including clinical applications in implantable and interventional cardiac therapies.

In this work, we approach the problem by leveraging atomically thin electronic graphene tattoos as soft, tissue-imperceptible, and transparent bioelectronic interfaces (Fig. 1). *Ex vivo* tests in a mouse model demonstrate the capability of cardiac electrogram recording and rhythm capturing and pacing by GET-electrodes. Compatibility of the transparent GET-electrodes with optical mapping and optogenetics allows accurate measurement and control of cardiac electrophysiology (EP) critical to cardiac function. In vivo tests in a rat model using an array of GETs indicate the potential of high-density electrical mapping from multiple locations on a beating heart. The successful ventricular rhythm restoration during the atrioventricular block (AV block), a common arrhythmia, indicates the potential of treating arrhythmias using GET-electrodes that can withstand the dynamic motion of the *in vivo* beating heart. This proof-of-concept study supports the use of transparent and flexible GET-electrodes fabricated by a simple and cost-effective method for applications in cardiac transvenous EP studies and potentially different implantable anti-arrhythmia devices.

**Figure 1.**
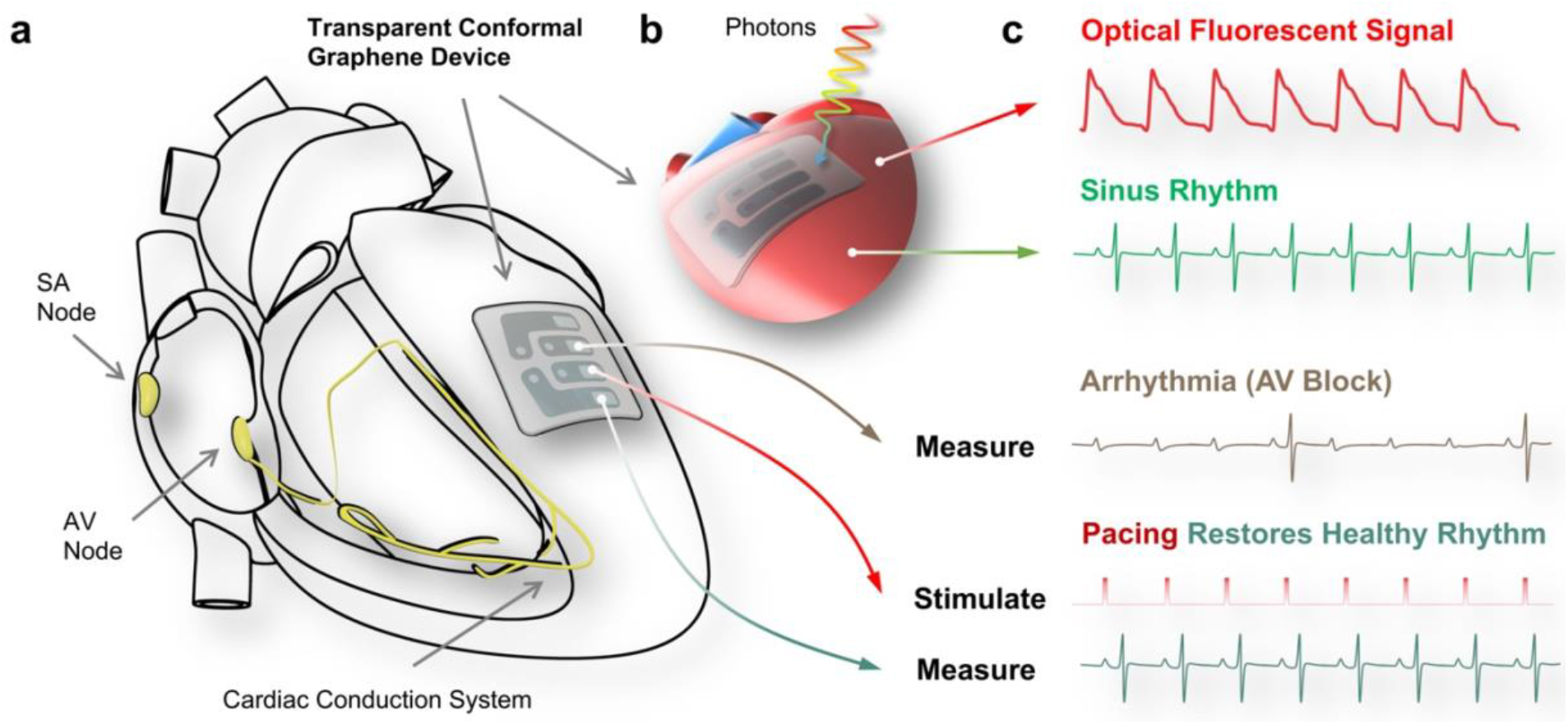
Schematic illustration of the graphene biointerface function. **a**, Illustration of a transparent, flexible, and tissue-conformable graphene electrode array placed on a heart that is also transparent and compatible with optical studies (**b**). **c**, Illustrative signals compatible with, recorded by the graphene electrode, showing the capacity to both record and stimulate the cardiac tissue, restoring healthy cardiac rhythms of the arrhythmia-affected tissues. SA node, sinoatrial node; AV node, atrioventricular node; AV block, atrioventricular block.

## Results & Discussion

### Graphene cardiac tattoos

The graphene biointerface technology reported here is derived from the previously introduced GETs^40,42,43^, additionally supported with two layers of ultrathin elastomer (Fig. 2). The GETs are made of atomically thin chemical vapor deposition grown graphene. The top layer of the elastomer features an electrode opening with a diameter varying between 1 and 3 mm. An ultrathin (10 μm) adhesive, elastic, and conductive gold tape was leveraged as the hybrid interconnect: soft enough for contacting graphene yet robust enough for external connection with measuring and stimulating electronics. Furthermore, to ensure that graphene does not affect biological processes *in vivo* or *ex vivo*, the GETs were modified with micrometer-sized holes (Fig. S2), allowing for efficient water transport. This feature also allows us to create robust electrical contact with the GETs. Besides contacting the GET from the graphene sides, we can actually build efficient contacts from the PMMA-side since the holes are embossed, and parts of the graphene are folded to the backplane (Fig. S2)^44^. Based on the knowledge from our previous works^40,44^, we decided to use 3-layer-GETs as they provide the most reproducible qualities, including the lowest sheet resistance and lower interface impedance.

**Figure 2.**
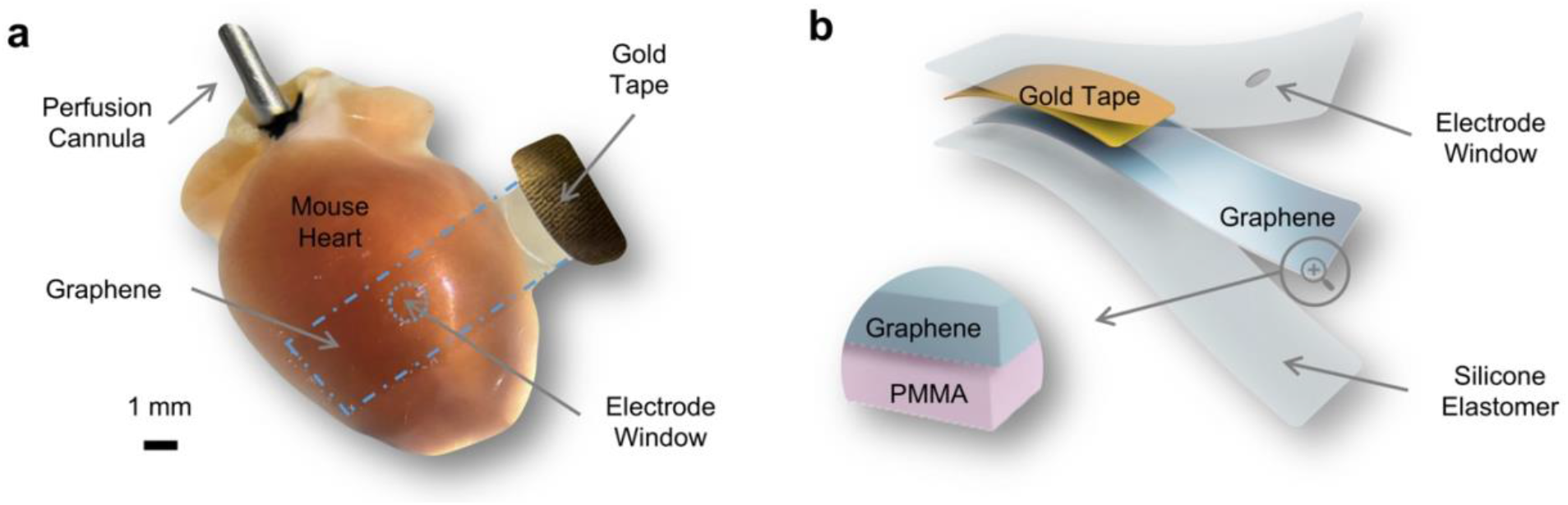
Soft and transparent graphene electrode. **a**, Optical photograph of a graphene electrode on a mouse heart. The graphene and the electrode window are highlighted with blue dashed lines. **b**, 3D schematic of the electrode featuring atomically thin and transparent graphene (blue) supported by 200 nm thick PMMA (pink) and encapsulated by silicone elastomer (light gray). The gold tape (yellow) serves as the interconnect between graphene and measuring and stimulating electronics. The diameter of the electrode window can be configured by using biopsy punches of various sizes (1-3 mm). PMMA, poly(methyl methacrylate).

A simple method (Fig. S3) was developed to produce GET-electrodes for a direct electrical interface with the heart (Fig. 2). First, poly(vinyl alcohol) (PVA) (10%, g/ml) is spin-coated (100 rpm, 40 sec, cure at room temperature for 48 h) on top of a glass slide, and acting as a temporary supporting layer that eases the delamination of the electrode from the glass slide. Then, Ecoflex silicone is spin-coated (100 rpm, 60 sec, cure at room temperature for 48 h), acting as a flexible encapsulating layer. Different electrode windows (*i.e.*, the exposure area for graphene contact with tissue) patterns are defined on the PVA and silicone layer with biopsy punches of various sizes (diameter 1-3 mm). The prefabricated adhesive gold tape is placed onto the flexible base silicone layer and used as interconnect. Then, the graphene tattoo is placed in such a way that one of its ends covers the electrode window, and the other end contacts the gold tape. The device is then encapsulated (spin-coated as aforementioned) by one more layer of Ecoflex silicone. The lightweight and soft GET-electrode could easily conform to the curvature of small anatomical structures of a mouse heart (Fig. 2a).

### Electrical and electrochemical characterization of graphene electrodes

To ensure reliability *ex vivo* and *in vivo*, the GET cardiac patches were first tested for their electrochemical performance. Large surface area GETs (typically 15 to 18 mm^2^) made for initial tests came in three configurations: monolayer graphene (1L), bilayer graphene (2L), and trilayer graphene (3L, see Methods for fabrication details). Chemical vapor deposition-grown graphene allows us to control the quality and performance of the devices over a large scale. Raman spectra of the 1L, 2L, and 3L GETs (Fig. S4) confirmed uniform high-quality graphene regardless of the multilayer stacking. Fig. 3a shows a clear trend in decreasing the electrochemical impedance of the devices with a larger number of graphene layers in the stack, similar to the trends when GETs were used as skin electrodes^40,44^.

**Figure 3.**
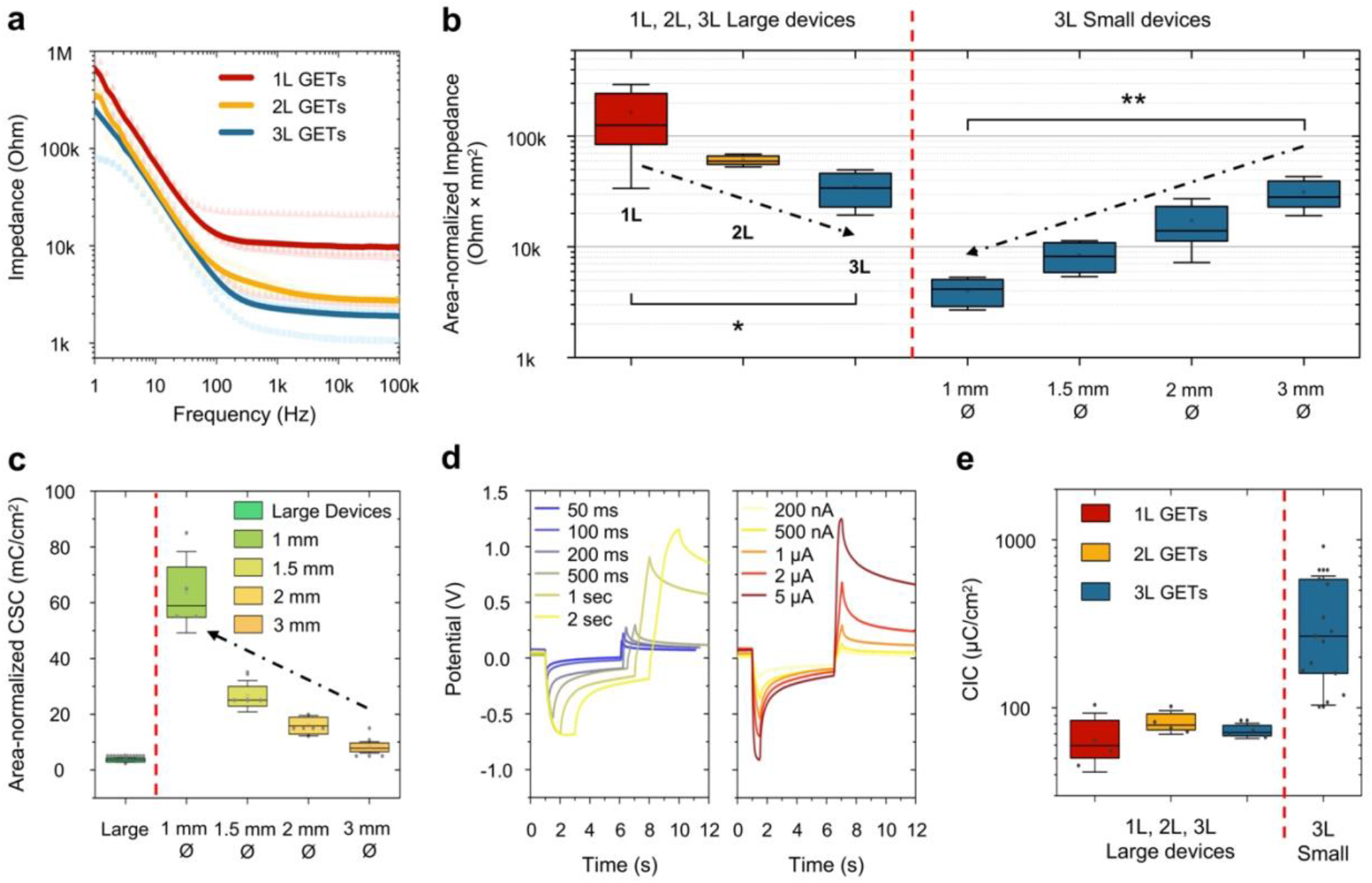
Electrochemical Performance of the GET arrays. **a**, Bode plot of the electrochemical impedance (plot on a logarithmic scale) of the large graphene tattoos, made of 1L (red), 2L (yellow), and 3L (blue) graphene (n=4 for each). Shaded symbols represent individual devices, and solid lines represent their average. **b**, Area-normalized impedance (plot on a logarithmic scale) of large GETs with 1L, 2L, and 3L configurations and small GETs with 1mm, 1.5mm, 2mm, and 3mm electrode diameters. For both large devices with different layers (i.e., left to the red dashed line) and 3 layers devices with different opening areas (i.e., right to the red dashed line), comparison was calculated with a nonparametric Kruskal-Wallis test in conjunction with Dunn’s multiple comparison test at a significance level of p < 0.05. n = 4 devices for each group. P-value = 0.024 (1L vs. 3L). P-value = 0.005 (1mm vs. 3mm). **c**, Area-normalized charge storage capacity of the large and small GETs of different electrode opening dimensions (large n=12, 1mm n=4, 1.5mm n=4, 2mm n=4, 3mm n=4). **d**, The voltage transients for various current widths (blue-to-yellow) with fixed amplitude and a set of different transients for various amplitudes (yellow to red) with fixed width. **e**, The area-normalized charge injection capacity (plot on a logarithmic scale) for GETs of different dimensions and number of layers. The box represents 25% and 75% with a mean, and the outliers are ±SD (1L large device n=4, 2L large device n=4, 3L large device n=4, 3L small device n=15).

On average, the 3L-GETs have an interface impedance of 2.5±0.7 kOhm @ 1kHz (Fig. S5) or area-normalized impedance of 345±151 Ohm×cm^2^ @ 1kHz (Fig. 3b). Since impedance raises with reduced electrode area, it is important to always report area-normalized impedance, in the Ohm×cm^2^ dimensions. Interestingly, the small devices with electrode openings of 1mm, 1.5mm, 2mm, and 3mm in diameter show a downward trend of reducing normalized impedance with smaller electrode openings, suggesting that smaller electrodes are more efficient (Fig. 3b and Table. S6). We hypothesize that the efficiency originates in the edge/surface ratio of the microelectrodes, with smaller electrodes having a greater ratio. A similar effect is reported in earlier works^45^. As the electrode size decreases, the reactants participating in the electrochemical reactions on the electrode surface diffuse from a planar to a hemispherical diffusion profile. The impedance change also transforms from an approximately r-2 scaling to an r-1 scaling^46^. The uneven current density distribution on the electrode surface described in Newman’s work^47^ also leads to the deviation in impedance scaling: the current density tends to be higher at the edges in both large and small disk electrodes. Therefore, both perimeter and area determine impedance scaling behavior. These results also explain the “edge effect,” which is increasing the electrode edge can reduce electrode impedance, leading to porous or array-shaped electrodes showing reduced impedance compared to planar electrodes of similar regions^48^.

The average value of area-normalized impedance for 1mm opening 3L GETs is 40±13 Ohm×cm^2^ at 1 kHz, which is on par and even exceeds the value of highly conducting PtTe2 and gold e-tattoos. The GETs exhibited outstanding stability and no degradation of the properties over 10 days of room temperature PBS exposure (Fig. S6 and Table. S5). To move forward with the electrochemical characterization of the cardiac graphene tattoos, we characterized their electrochemical stability and determined their so-called water window^49^. As well-known from previous works, graphene has a very wide water window, which we corroborated in this work. We have been able to safely perform cyclic voltammetry (CV) (Fig. S7) measurements in the range of applied potential between −0.9V and +1.2V, which defines the graphene water window. Furthermore, we have found no significant dependency of the water window per electrode opening area or per number of graphene layers (Fig. S7). The CV measurements, performed at a 5 mV/sec rate, have been used to estimate another essential figure-of-merit of microelectrodes, their charge storage capacity (CSC). Similar to the case of interface impedance, the area-normalized values of CSC are much larger for the smaller electrodes (Fig. 3c). For large devices, the average CSC is estimated in the range of 2-5 mC/cm^2^. At the same time, the value is almost an order of magnitude higher, 63.7±14.6 mC/cm^2^, for smaller electrodes, with ~1mm diameter opening (Table. S7). Finally, the essential value for effective stimulation of electrogenic tissue is the charge injection capacity (CIC) of the electrodes. For this purpose, the charge-balanced measurements with constant-current pulses were applied to the electrodes. The resulting voltage transient for various pulse amplitudes and pulse widths are shown in Fig. 3d. Extracting the CIC capacity gives us a somewhat similar performance for 1L, 2L, and 3L large GETs, in the range of 60-90 μC/cm^2^, with the values drastically increasing for smaller devices (Fig. 3e), reaching up to 704±144 μC/cm^2^ for electrodes with ~1mm diameter opening (Table. S8). One can note a slight upwards trend with increasing the number of layers, suggesting that multilayer graphene helps in out-of-plane charge injection. The recorded properties for large devices are in good accordance with other works, while the superior performance measured for small GET-arrays is unprecedented. Compared to the state-of-the-art (Table. S4), the graphene electrodes are the thinnest made biointerfaces with the lowest (for graphene) interface impedance. Only a handful of works, mainly utilizing PEDOT and porous metals, can compete with the GETs. The same trend can be found for CSC and CIC except for composite multi-coated electrodes (*e.g.*, CNTs + porous Pt).

### *Ex vivo* cardiac electrophysiology sensing and actuating characterization

Upon completion of electrochemical characterization of the GET-electrodes, they were applied for monitoring cardiac electrical activity in an *ex vivo* Langendorff-perfused mouse heart model where the electrogram recorded by graphene (gEG) was compared with the traditional far-field ECG simultaneously recorded (with commercial needle electrodes made of Ag/AgCl, MLA1203, ADInstruments) in the perfusion bath (Fig. 4a). The ECG was recorded from three sensing electrodes in the ECG lead II position. The gEG was recorded from a 2-sensing electrodes setup. The unipolar GET-electrodes (1, 1.5, and 2 mm electrode window size) were used as the positive (+) electrode paired with another negative (-) electrode (made of Ag/AgCl). The simultaneously recorded ECG and gEG (Fig. 4b-c) showed a good temporal correlation between R waves and elapsed time between two successive R waves (RR interval, inversely related to heart rate) (Fig. 4d). Because the GET-electrodes were placed on top of the left ventricle with electrode windows close to the apex of the heart, signals recorded were local electrogram, and the P waves were absent from gEG signals (Fig. 4c), present in ECG, which characterizes whole heart electrophysiology. The signal-to-noise ratio (SNR) was compared (Fig. 4e) for gEG recorded from unipolar GET-electrodes with various window sizes. We found that gEG of 1 mm samples showed comparable performance to ECG. At the same time, there was a statistically significant decrease in the SNR of gEG recordings between the 1 mm and 2 mm samples. Only in the 2 mm group the gEG SNR was significantly lower than the control ECG SNR. This experimental data correlates with the data shown in Fig. 3b-c, where smaller diameter electrode arrays have featured superior interface impedance, charge storage, and charge injection capacities.

**Figure 4.**
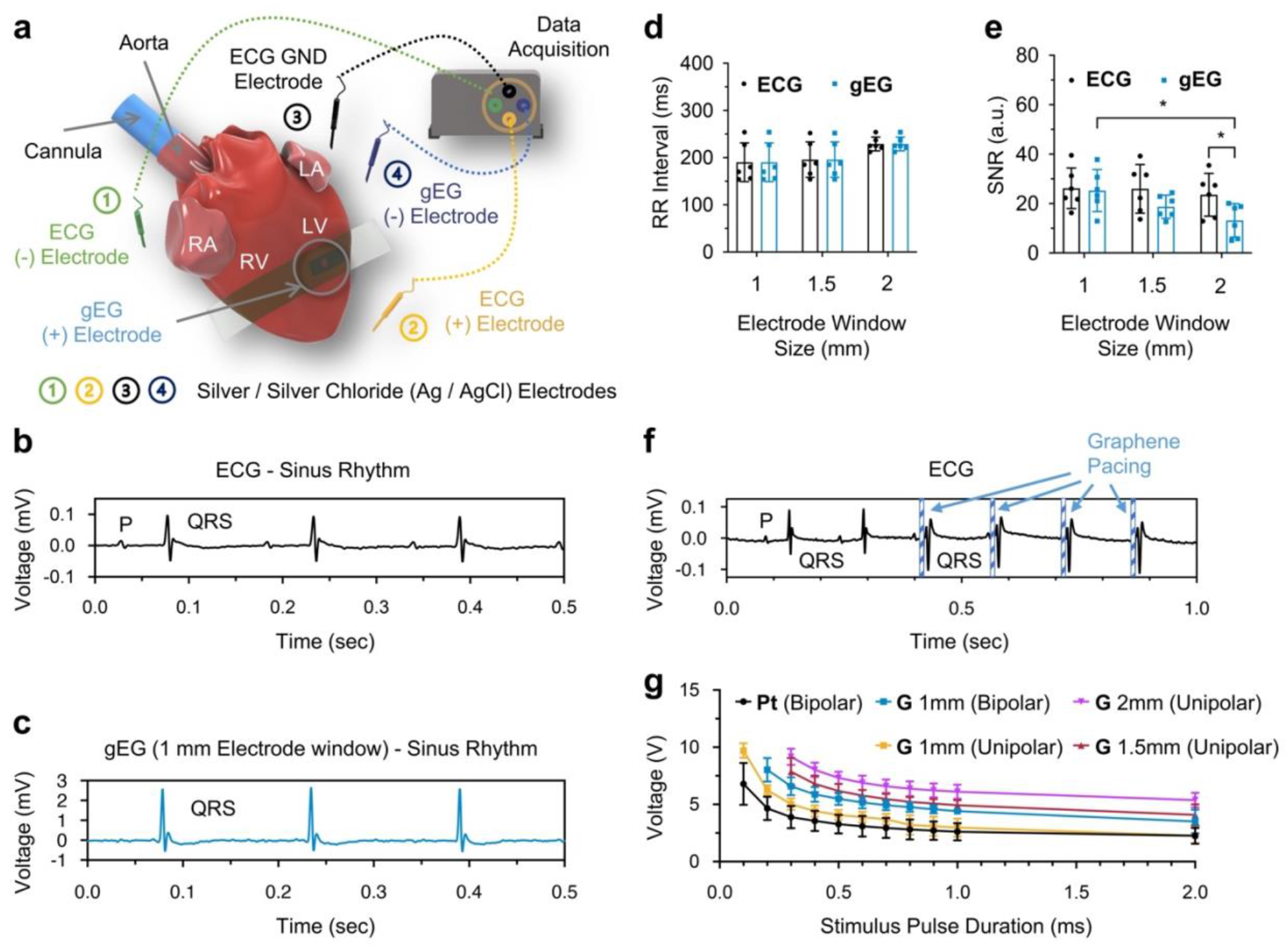
Ex vivo cardiac electrophysiology sensing and pacing with GETs. **a,** Schematic of the ex vivo study setup. The GET-electrode was placed on a Langendorff-perfused mouse heart at the anterior side of the left ventricular surface. Far-field pseudo-ECG (ECG) was recorded using a 3-sensing electrodes setup. The graphene electrogram (gEG, blue) was recorded using a 2-sensing electrodes setup. **b-c,** Simultaneously recorded ECG **(b**, black) and gEG **(c**, blue) at natural sinus rhythm (heart rate = 385 beats per minute). **d,** Sinus rhythm RR interval (i.e., heart rate) calculated from ECG and gEG. n = 6 hearts for each group. **e,** Signal-to-noise ratio (SNR) comparison. n = 6 hearts for each group. All the graphene electrodes were in a unipolar window setup but with different window diameters. The intragroup comparison (i.e., ECG vs. gEG) was calculated with a nonparametric Wilcoxon test at a significance level of p < 0.05. The intergroup (i.e., 1- vs. 1.5- vs. 2 mm gEG) comparison was calculated with a nonparametric Kruskal-Wallis test in conjunction with Dunn’s multiple comparison test at a significance level of p < 0.05. P-value = 0.0242 (1 mm gEG vs. 2 mm gEG). P-value = 0.0312 (2 mm ECG vs. 2 mm gEG). **f,** Cardiac pacing (pulse duration = 2 ms, pulse amplitude = 2.5 V, cycle length = 150 ms) was achieved by using the GET-electrode. **g,** The ex vivo pacing strength-duration curve. n = 6 hearts for each group. Data are presented with error bars as mean ± SD. RA, right atrium; LA, left atrium; RV, right ventricle; LV, left ventricle; Pt, platinum; G, graphene.

Cardiac actuating was also achieved by connecting the GET-electrodes to the pulse generator (PowerLab 26T, ADInstruments). Besides the unipolar GET-electrodes, bipolar GET-electrode (two 1 mm electrode windows, window pitch distance of 2 mm) were also tested. In the unipolar actuating mode, the GET-electrode was used as the positive (+) electrode paired with another negative (-) electrode (made of Ag/AgCl). In the bipolar mode, the two graphene electrodes were used as (+) and (-) electrodes, respectively. Successful ventricular pacing (i.e., the capture of heart rhythm) resulting in a faster heart rate than sinus rhythm was confirmed by ECG morphology in which each pacing artifact was constantly followed by a wide QRS complex with tall and broad T-wave, as commonly observed in clinical pacing setup (Fig. 4f). The pacing strength-duration curve relates pacing pulse duration to pacing threshold (expressed in voltage / current / energy) required to elicit excitation (*i.e.*, cardiomyocytes stimulation). Pacing strength-duration curve can be used (1) to evaluate the electrical characteristics of different electrodes (e.g., shape, material) and (2) to provide information on choosing optimal stimulation intensity and stimulation duration setup that maximizes the battery life of implantable pacemakers^50^. Here the *ex vivo* pacing strength-duration curve was characterized for various GET-electrodes along with a custom bipolar platinum (A-M Systems, catalog number 778000, diameter = 0.127 mm) electrode that served as the reference since it is the electrode our lab always uses for optical mapping studies (Fig. 4g, Table. S9). The choices of stimulating pulses duration included both animal research related values (*e.g.*, 2 ms for optical mapping studies) and clinically relevant values (*i.e.*, from 0.1 to 0.5 ms)^51^. With a decrease in pacing duration there was an increase of voltage threshold as expected. At all pulse duration, there was a trend of having higher voltage threshold with bigger electrode surface area (surface area of bipolar platinum < 1 mm unipolar GET < 1 mm bipolar GET < 1.5 mm unipolar GET < 2 mm unipolar GET). It has been reported that electrodes of similar configuration and the same material but of different surface areas require the same energy density to produce a cardiac contraction^52^. Therefore, higher total amount of voltage is required for graphene electrodes with a bigger surface area to excite cardiomyocytes. The 1 mm unipolar GET electrode, which had the lowest pacing threshold compared with other GET electrodes and had no significant difference in voltage threshold compared with platinum electrode (at 2 ms pulse duration), was therefore incorporated into the subsequent optical mapping study.

### Validation of GET-electrodes with optical mapping studies

Graphene is an optically transparent material reported to make transparent electrodes^28,38^. Cardiac optical mapping is a fluorescent imaging technique frequently used in cardiac physiology to study the excitation-contraction coupling between transmembrane potential and calcium handling using reporters that emit fluorescence upon excitation illumination^11^. Here GET-electrodes were incorporated into optical mapping studies where light transmission is critically important. It is common to use metal electrodes such as platinum to stimulate the heart during experiments. However, metal electrodes optically block the partial area of the tissue preventing optical recordings (Fig. S8). Depending on the electrode size, it may be difficult (*e.g.*, low optical signal amplitude, distorted signal morphology) or even impossible to analyze signals from pixels in the region of interest. By using the transparent GET-electrode, those issues are now resolved.

Here we measured and compared the cardiac restitution properties recorded (Fig. 5a) from optical signals (*i.e.*, action potentials and intracellular calcium transients) during pacing by a unipolar GET-electrode (1 mm window size) and a custom bipolar platinum electrode. The GET-electrode, which was placed on the anterior side of the left ventricular surface of the heart, could barely be seen in a bright-field image captured by the camera (Fig. 5b). The activation map showed the expected anisotropic propagation of the transmembrane potential originating from the site of cardiac pacing by GET-electrode throughout the ventricular myocardium (Fig. 5c). Clear representative optical signals from different heart locations, including where the GET-electrode was aligned well with the simultaneously recorded ECG (Fig. 5d). The wide and high-amplitude QRS complex immediately after each pacing artifact indicated successful ventricular capture by the GET-electrode. No statistical difference (Fig. 5e) was found between the platinum and graphene group of action potential duration 80 (APD80), calcium transient decay constant (Ca Tau), as well as longitudinal and transverse conduction velocity (CVL, CVT). Taken together, these tests indicate that the flexible and transparent GET-electrode is readily applicable to optical mapping studies with high efficacy in cardiac pacing and no harm to the precise measurement of cardiac restitution properties. Besides, we tested the GET-electrodes for potential cross-connection and insulation performance. As one can see from the control experiments (Fig. S10), samples without graphene or passivation opening yield no ECG recording, ensuring that the signal originates specifically from graphene.

**Figure 5.**
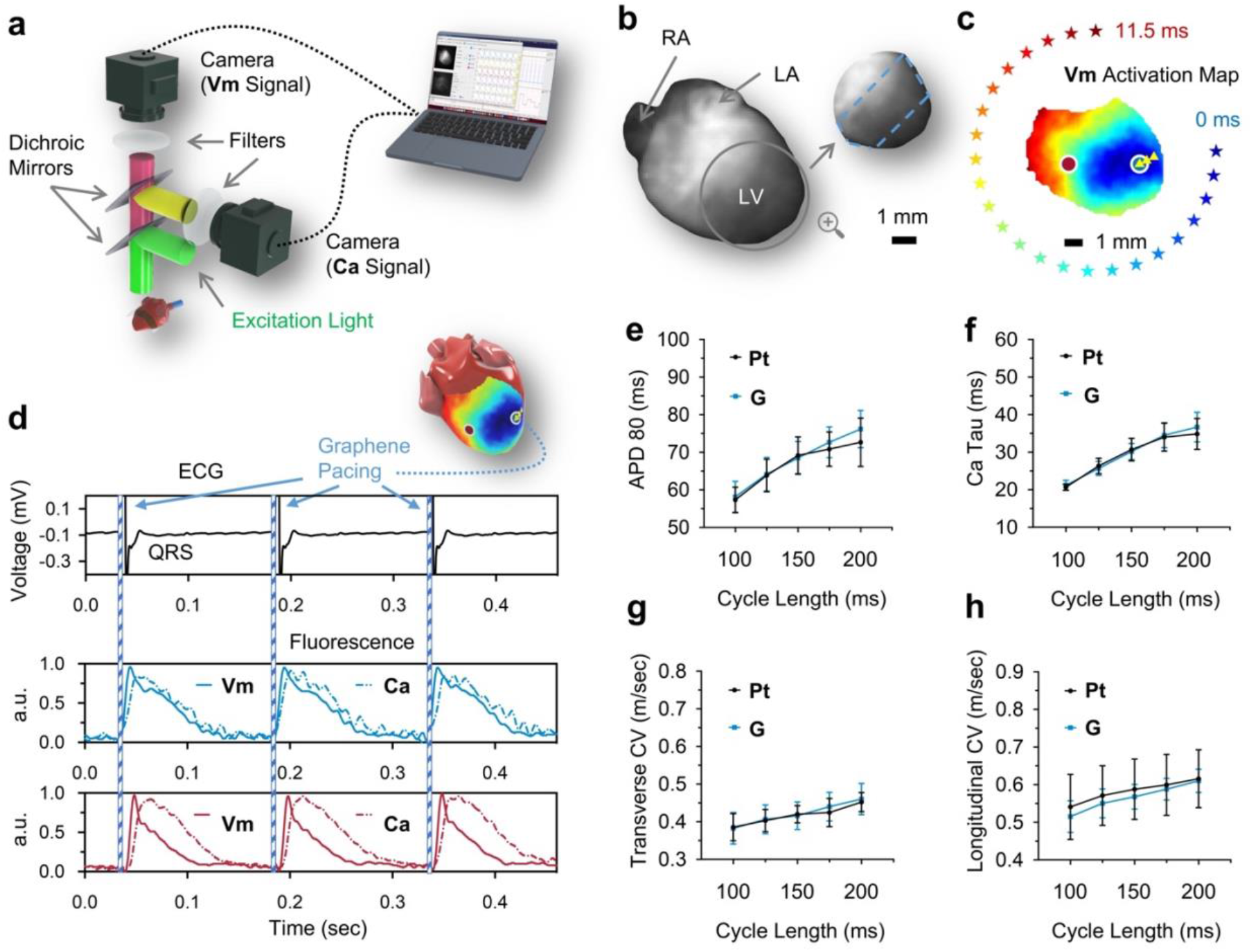
Combination of graphene electrodes with optical mapping studies. **a,** Epi-illumination optical mapping system configuration and the light path for Vm and Ca signals. The Langendorff perfused heart was stained with voltage (RH237) and calcium (Rhod-2 AM) sensitive dyes. Emission fluorescence, separated by wavelength using dichroic mirrors and optical filters, was recorded using two CMOS cameras with 100 × 100 pixels resolution (ULTIMA-L, SciMedia) at 2 kHz sampling frequency. **b**, Optical mapping camera view of a mouse heart with a magnified view of the heart’s left ventricle on which a transparent unipolar (1 mm window size) GET-electrode was placed. The edge of the graphene is highlighted with blue dashed lines. **c,** Activation map of the transmembrane potential during electrical stimulation by the GET-electrode. The yellow triple-triangle symbol marks the location of the electrode window of the GET-electrode. The electrical stimulation is performed right at the spot of the electrode opening (blue circle). The blue and red circles mark the location from which representative traces of Vm and Ca (solid and dashed traces, respectively) were recorded (d). **d,** Simultaneously recorded ECG and optical signals. The wide QRS complex aligned well with optical signals, representing successful ventricular activation upon cardiac pacing (pacing pulse duration= 2 ms, pulse amplitude = 4.5 V, cycle length = 150 ms). **e-h,** Summary restitution properties of the four parameters measured simultaneously by optical mapping. Nonlinear regression analysis was performed on APD80 (**e**), Ca Tau **f**), CVT (**g**), and CVL (**h**) using the least-squares regression fitting method. An exponential curve Y = Y_M_ – (Y_M_ – Y_0_) * e^-K*X^ was used. Goodness-of-fit was determined by r-squared value > 0.5. The null hypothesis, whether one curve adequately fitted all data sets, was tested with the extra sum-of-squares F test at a significance level of p < 0.05. **e-h,** n = 6 hearts for both platinum and graphene groups. Data were presented with error bars as mean ± SD. RA, right atrium; LA, left atrium; LV, left ventricle; Pt, platinum; G, graphene.

### GET-electrodes electrical recording with optogenetic pacing studies

GET-electrodes were incorporated into optogenetics studies to evaluate the general utility of graphene electrodes for simultaneous electrical sensing and optical pacing (Fig. 6). Transgenic mice expressing light excitable opsins, channelrhodopsin-2 (ChR2)^12^, in cardiomyocytes were used. Cardiac optical pacing works as the opening of ChR2 upon blue light (470 nm) illumination allows cations (e.g., Na^+^, K^+^, Ca^2+^) to enter cardiomyocytes which in turn elevates the transmembrane potential to elicit action potentials at a cell level and ECG at a whole heart level. Similar to optical mapping studies, the transmission of light is critically important.

**Figure 6.**
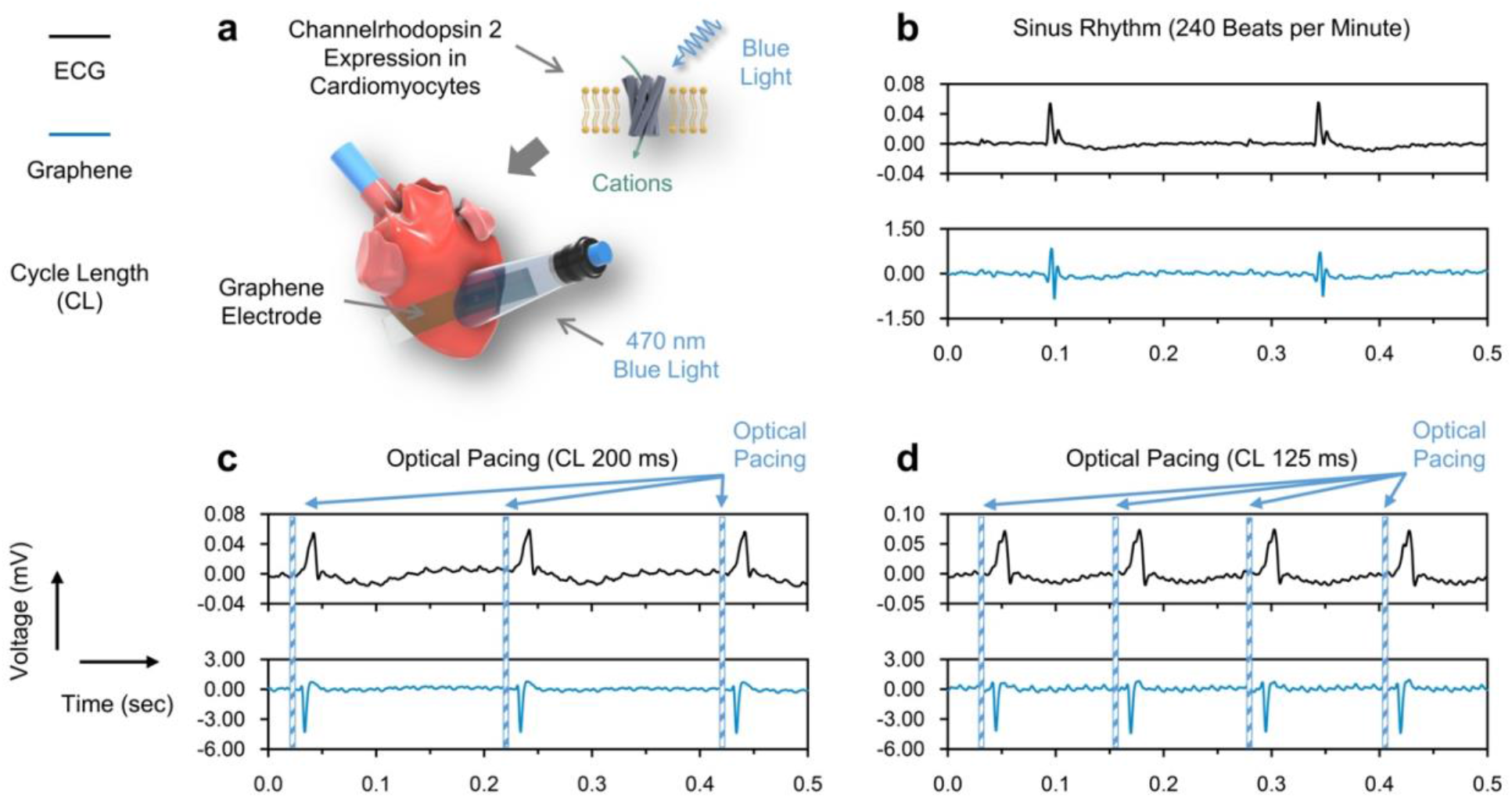
Combination of graphene electrodes with optogenetics studies. **a,** Schematic of the ex vivo study setup. Optical pacing was achieved using mouse hearts with channelrhodopsin 2 (ChR2) expression in cardiomyocytes. Cations enter cardiomyocytes after the opening of ChR2 upon blue light illumination and elicit action potential at a cell level and ECG at a whole heart level. **b**, Simultaneously recorded ECG (top, black) and gEG (bottom, blue) at natural sinus rhythm (heart rate = 240 beats per minute). **c,d,** Simultaneously recorded ECG and gEG upon optical pacing at different cycle length (200 ms in **c** and 125 ms in **d**).

The GET-electrode conformably contacted the left ventricle of the mouse heart (Fig. 6a). Photons from an external blue LED (470 nm) passed through the transparent electrode and optically paced the underneath cardiomyocytes. The simultaneously recorded ECG and gEG showed natural cardiac bioelectrical activates (Fig. 6b) as well as signals during optogenetic pacing at cycle length of 200 ms (Fig. 6c) and 125 ms (Fig. 6d). Results from the electrophysiology, optical mapping, and optogenetics studies together indicate that the lightweight and transparent GET-electrodes allow seamless interfacing with cardiac tissue yet efficient light transmission for precise and simultaneous optoelectronic recording and actuating of cardiac bioelectrical activities.

### GET-electrode-array for cardiac electrical mapping

Electrical mapping with the high spatial resolution is critical during cardiac EP studies required to guide ablation therapy of arrhythmias. For example, unsatisfactory spatial resolution can result in the incorrect location of drivers of atrial fibrillation (AF), which is the most common type of cardiac arrhythmia affecting millions of patients^53^. To prove the potential high spatial resolution cardiac electrical mapping ability of electrodes based on GETs, we fabricated a novel array of micropatterned GETs (mGETs) and showcased its sensing ability on an *in vivo* beating rat heart (Fig. 7a-c). Unlike earlier version GETs with a whole graphene layer supported by a layer of ultrathin (200 nm) PMMA, the mGETs (Fig. 7a) have two unique features. First, the layer of graphene was patterned into multiple feedlines and individually addressed, all being supported by a single PMMA substrate. Secondly, the mGET-arrays were additionally passivated on top with another layer of PMMA with openings only at the contact and electrode opening sites (like classic microelectrode arrays). Importantly, the PMMA-graphene-PMMA arrays do not have micro-holes in the structure in order to ensure passivation integrity and ability to record from only required passivation window openings. Meanwhile, this PMMA-graphene-PMMA stack resulted in thinnest even microelectrode array, with a thickness below 500 nm. Importantly, such devices can be fabricates using off-the-shelf, low-cost additive fabrication components (see Methods). Besides, the mGET arrays can be scaled up and high-density devices can be mass-produced like the classic graphene-based microelectrode arrays or field-effect transistors^38,54,55^.

**Figure 7.**
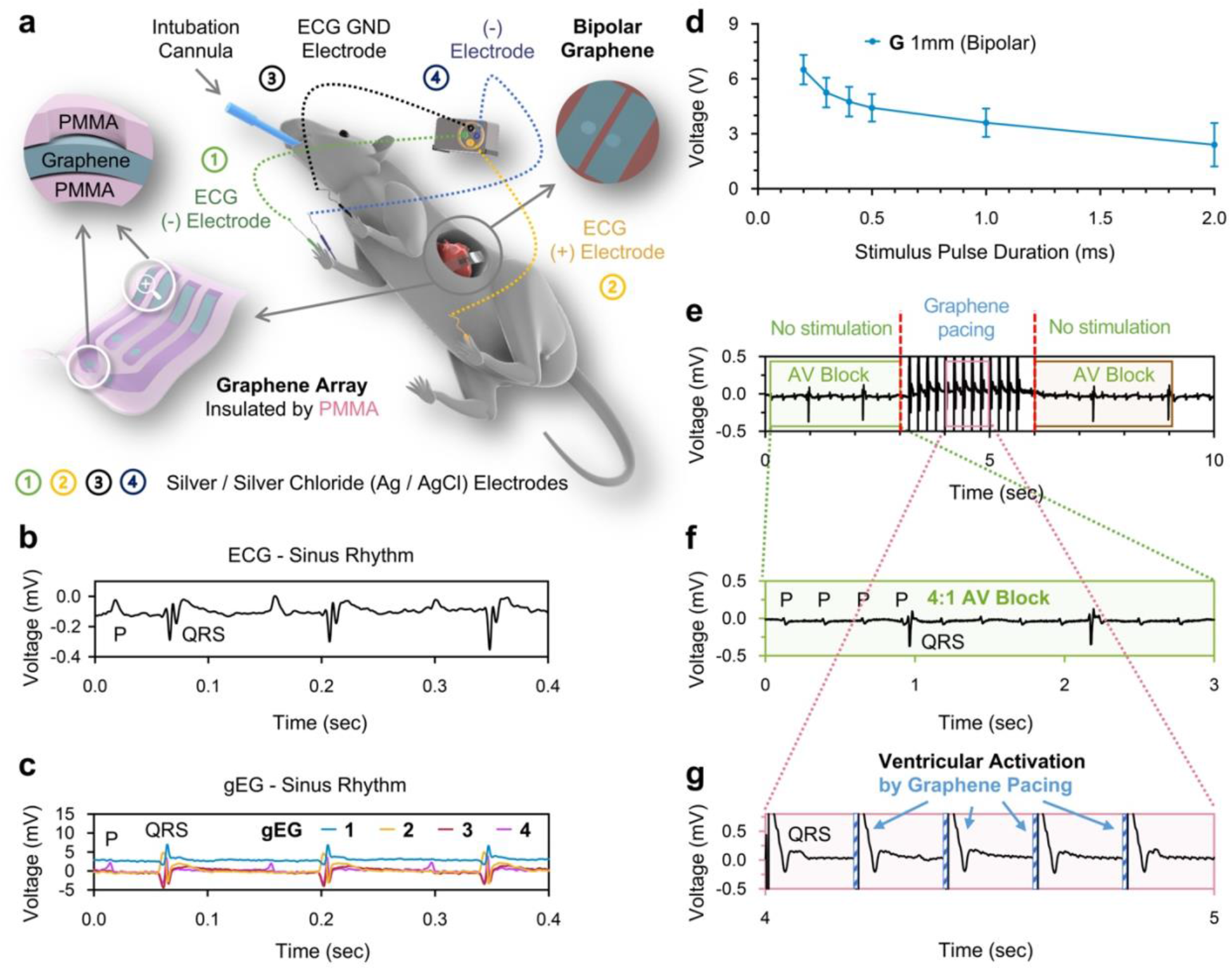
Cardiac electrical mapping and treatment of AV block in an in vivo rat model. **a,** Schematic of the in vivo study setup. ECG was recorded through commercial needle electrodes positioned in the Lead II configuration. Electrogram was recorded using the 2-by-2 mGET-array-electrode (the magnified view shows an electrode array of two PMMA layers highlighted with purple color and graphene in light blue). Electrode window size of each graphene feedline = 1 mm. The window pitch distance = 6 mm. Bipolar GET-electrode had 1 mm window size and 2 mm window pitch distance. The GET-array-electrode could simultaneously record gEG from both the right and left ventricles of the heart. **b-c,** Simultaneously recorded ECG (**b**) and four gEGs from four electrodes within one array (**c**) at natural sinus rhythm (heart rate = 425 beats per minute). gEG-4 showcases the ability to record atrial electrical signals (i.e., P wave) when the graphene electrode is placed near the atrium. **d,** In vivo pacing strength-duration curve using bipolar GET-electrodes. In bipolar pacing mode, one and the other graphene electrodes were used as (+) and (-) electrodes, respectively, n = 6 hearts. **e-g,** In vivo ECG monitoring of a rat heart with 4:1 AV block before cardiac pacing (green box), regular ventricular activity during pacing (cycle length = 200 ms, pacing pulse duration = 2 ms, pulse amplitude = 2.5 V) (pink box), and AV block when pacing was stopped (brown box). Ventricular capture via electrical stimulation was observed as the wide QRS complex after each pacing artifact. G, graphene. PMMA, poly(methyl methacrylate).

The 2×2 mGET-arrays were placed on a rat heart to record electrical signals from both right and left ventricles (Fig. 7a-c). *In vivo* ECG was recorded throughout the period of the experiments (about 90 minutes) from sensing electrodes positioned in the lead II configuration. One can see prominent PQRST phases of a healthy cardiac wave. Because one graphene electrode (Fig. 7c, gEG-4) was placed close to the atrium, a clear P wave was observed. The simultaneously recorded ECG and gEG showed a good temporal correlation of R waves. Fig. S11 shows the four gEGs individually.

### Treating atrioventricular block (AV block) in an *in vivo* rat model

Fig. 7d shows the *in vivo* pacing strength-duration curve of bipolar GET-electrodes. Stimulus pulse duration covered the clinically used values such as from 0.2 to 0.5 ms^51^. Fig. 7e-g and Supplementary Video 1 show the application of the bipolar GET-electrode to the treatment of AV block. AV block was induced by fast pacing (cycle length 100ms) at the left ventricle after intraperitoneal injection of 120 mg/kg caffeine and 60 mg/kg dobutamine. When no pacing treatment was applied, the rat heart exhibited continuous 4:1 AV block (Fig. 7e-f). Upon ventricular pacing using the GET-electrode (pacing heart rate at 300 BPM), successful capture with clear pacing spikes and widen morphologies of the QRS complex was observed (Fig. 7g). The ventricular beating was also apparent during pacing compared with almost silent contraction during AV block (Supplementary Video 1). When pacing was stopped, the heart rhythm returned to AV block (Fig .7). Overall, these *in vivo* tests demonstrate that the flexible mGET-arrays can conform reliably to the curvilinear topography of a beating heart (90 minutes of study time × 350~380 beats per minute = over 30,000 beats) without hampering cardiac contraction to achieve in vivo rhythm sensing and modulation, which potentially can be applied to clinically relevant scenarios like ventricular rhythm resuscitation after bradycardia or AV block. This is the first-time graphene electrodes have been used to successfully treat a life-threatening heart rhythm disorder.

## Conclusions

Bioelectronics has revolutionized cardiology and cardiac physiology by enabling electrocardiography, arrhythmia diagnostics, anti-arrhythmia ablation, and implantable device therapies. The diagnosis and treatment of cardiac diseases (*i.e.*, arrhythmias, myocardial infarction, and heart failure) have also been facilitated since the arrival and evolution of essential tools such as cardiac pacemakers and defibrillators. In recent years, the conventional bulky and rigid cardiac devices and catheters show a clear trend of becoming smaller, softer, portable, and multifunctional by engineering better materials or designing more sophisticated fabrication methods. Yet, there is still no technology capable of seamless integration with the perpetually undulating cardiac tissue that would robustly enable electrical sensing and actuation of the tissue properties.

In this study, we report on a distinct class of soft and transparent bioelectronics fabricated using graphene electronic tattoos. The sub-micrometer thickness and softness of graphene *in vivo* biointerfaces make them superior for seamless coupling with the cardiac tissue, yielding unique elements of soft bioelectronics that conform well to the heart without constraining or altering its natural motions. Using off-the-shelf fabrication tools allows us to easily change and personalize the interfaces, depending on the application of the future studies. Electrochemical properties, including interface impedance, charge storage capacity, and charge injection capacity of the reported 500 nm thick PMMA-supported and passivated graphene biointerfaces are superior to most emerging technologies (Tables. S2-4). While sensing and actuating cardiac tissue, leveraging transparency of graphene biointerfaces, we can simultaneously perform optocardiography, *i.e.*, using light to sense and stimulate electrical or metabolic dynamics of the heart, as demonstrated here *via* a set of systematic *ex vivo* and *in vivo* animal studies. Most importantly, we show that the graphene biointerfaces can effectively diagnose and treat a common clinical arrhythmia, AV block, in a rat model, signifying their superiority for future high-density electrical pace-mapping *in vivo*, and leading to advanced diagnosis and treatment of cardiac arrhythmias.

Furthermore, there is still plenty of room for improvement over this proof-of-concept work. First, the efficacy of epicardial electrical recording and stimulation might be hampered in the presence of cardiac fat tissue. When translating into larger mammals, however, endocardial sensing and actuation will be preferential, and it typically does not contain much fat. Finally, for the long-term implantation scenario, tissue growth over devices is an important issue to be premeditated. Although we anticipate little to none toxicity from the substrate-supported graphene, additional adhesive and anti-inflammatory coatings^56^ would be helpful.

## Methods

### Preparation of three-layer graphene tattoo

Graphene requires adequate protection and support, provided using the ultrathin layer of PMMA. High-quality monolayer graphene is grown *via* the CVD method on a copper foil (Grolltex), fixed the graphene/copper piece on a Si wafer by Kapton tape, and then a layer of PMMA (950 A4) is spin-coated at 2500rpm for 60 seconds, followed by 200°C bake on the hotplate for 20 min. Copper was removed by placing the PMMA/graphene/copper in 0.1 M (NH_4_)_2_S_2_O_8_ solution for 8-12h. The floating PMMA-graphene was then picked up by a Si wafer (must be larger than the piece of PMMA/graphene flake) and transferred into DI water in three stages (5min, 5min, and 30min) to remove any residual copper etchant solution. The PMMA/graphene piece was then picked up by a new piece of clean 1L graphene/copper. PMMA-graphene-graphene-copper was air-dried overnight at room temperature in a hood. Two-layer graphene was produced by annealing the PMMA-graphene-graphene-copper on a 200°C hot plate. The copper foil of the 2-layer graphene was also removed by putting it in 0.1 M (NH4)2S2O8 solution for 8-12h. 3-layer graphene on copper foil was fabricated by repeating the preceding steps. Using two sheets of tattoo paper (Silhouette Cameo), transfer PMMA/graphene and flip to have an arrangement with graphene facing up^40^.

### Cut the graphene tattoo into 15mm*3mm strips

The graphene tattoo was attached to Silhouette Cameo cutting mat using Kapton tape. A Cameo plotter was used to cut the graphene tattoo into 15mm*3mm strips (Cut Through graphene and tattoo paper: Depth 2, Pressure 25; Scoring Surface, only cut graphene: Depth 1, Pressure 15). To get a clean designed shape, immerse the graphene tattoo in water for about 30 seconds, use tweezers to remove unwanted parts, and separate each graphene strip.

### Preparation of graphene electrode

(Supplementary Fig. 2) 10% (g/ml) poly(vinyl alcohol) (PVA) (Millipore Sigma, catalog number 341584) was spin-coated (100 rpm) for 40 sec onto a glass substrate and cured at room temperature (20~22 °C) for 48 hours. The water-soluble PVA layer served as a temporary supporting layer to ease the delamination of the encapsulation layer (made from Ecoflex silicones) from the glass substrate. Ecoflex™ 00-30 (Smooth-On) silicones were prepared by mixing Ecoflex monomer and curing agent (volume-to-volume, v/v = 1:1). Vacuum degassing was applied for 5~10 minutes until bubbles were cleared. The liquid silicones were spin-coated (100 rpm) for 60 sec on top of the PVA layer and cured at room temperature for 48 hours. The electrode window with a desired diameter and pattern was created by cutting through the silicone encapsulation layer with biopsy punches. The unipolar pattern had an electrode window size of 1-, 1.5-, and 2 mm. The bipolar pattern had two 1 mm electrode windows with a pitch distance of 2 mm. The 2 × 2 array pattern had four 1 mm electrode windows with a pitch distance of 6 mm. A gold tape worked as the interconnect and was gently placed on top of the silicone encapsulation layer. And the graphene tattoo was gently placed in such a way that one of its ends covered the electrode window, and the other end contacted the gold tape. Another layer of silicone encapsulation was prepared as aforementioned. Just before use, the graphene electrode was gently peeled off from the glass substrate, and the PVA layer was washed off by 1X PBS solution.

### Preparation of micropatterned graphene electrode arrays

In order to prepare the 3L-GET arrays, an identical strategy described above of creating 3L-GETs is used, with the only exception that PMMA is annealed at only 150°C. When 3L graphene is then stacked on the copper foil, it is soaked in acetone, dissolving the PMMA. The 3L graphene on copper foil is then fixed on a dummy silicon wafer, spin-coated with a photoresist, exposed using plastic mask (with mm-large features printed using office printer), and exposed areas are treated with oxygen plasma to etch away graphene. The photoresist is then stripped off, and the structure is coated with another layer of PMMA, baked at 200°C, followed by copper etching. In parallel, a new piece of tattoo paper (Silhouette), after removing the top PVA layer, is coated with bare PMMA, baked at 200°C. Then, using Silhouette Cameo, and exactly same design used to print UV masks, the passivation openings (for electrodes and contacts) are cut-out from the PMMA layer. This piece will served at substrate to fish the graphene/PMMA piece. The transfer is performed by floating the graphene/PMMA piece, and using the PMMA/tattoo paper to fish the 3L graphene array up with precision and alignment. The two parts, graphene/PMMA and PMMA/tattoo paper are then dried overnight and annealed at 150°C, forming the thinnest to-date biointerface arrays.

### Electrical Impedance Characterization

The EIS, CV, CSC, and CIC characterization was performed on CEC Autolab TGSTAT 128N in a three-electrode system. Graphene was connected as the working electrode (WE), the platinum wire as the counter electrode (CE), and a 2.0mm diameter Ag/AgCl pellet as the reference electrode (RE), all immersed in PBS×1 solution. The graphene electrode was connected by gold tape and conductive silver epoxy. *For EIS measurements*, 10 mV sine waves at frequencies from 1 Hz to 100 kHz were applied. *For CV measurements*, the potential sweep from −0.9 to 1.2 V was applied, which is a water window, and the cycle curve was plotted to show the stabilized signal, with scan rates ranging from 5 to 1000 mV/s.

#### To estimate charge storage capacity (CSC)

the CV curve which at water window with a scan rate of 5 mV/s was used. The areas of cathodic (CSC_C_) and anodic (CSC_A_) were integrated and calculated the area difference (see Fig. S9a). The total CSC was obtained by subtracting the |CSC_C_-CSC_A_|, and normalizing per the exposed geometric area of the devices. *For charge injection capacity (CIC) measurements*, charge-balanced, constant-current pulses with a current range between 0.1μA to 10μA, cathode leading, 0.5s long phases, and 5s gap between phases were applied to the WE (return path through the CE). The resulting voltage transient was measured between the WE and RE. Gradually increase the pulse current until the working electrode voltage reaches the water window. The maximum negative polarization potential (Emc) was assumed to be the WE voltage 50 μs following the end of the cathodic phase, and the positive polarization potential was also 50 μs following the end of the anodic phase. The CIC was obtained by current at Emc multiplied by the time (which is 0.5s) divided by the exposed geometric area of the devices (see Fig. S9).

### Animals

The George Washington University Institutional Animal Care and Use Committee approved all experimental animal protocols. The procedures were by suggestions from the panel of Euthanasia of the American Veterinary Medical Association and the National Institutes of Health Guide for the Care and Use of Laboratory Animals. *Ex vivo* studies were performed on mouse (15~25 weeks old; C57BL/6J; Jackson Labs stock # 000664; male and female) hearts. For ex vivo optogenetic pacing studies, mice (15~25 weeks old) expressing channelrhodopsin 2 in cardiomyocytes were used. See “Method - Mouse model for optogenetics” for model creation details. *In vivo* studies were performed on Sprague-Dawley rats (15~20 weeks old; Hilltop Lab Animals; female)

### Langendorff perfusion of the heart for ex vivo studies

Mice were anesthetized by 3% (volume-to-volume, v/v) isoflurane inhalation until the animal no longer responded to a toe pinch. The cervical dislocation was quickly performed before thoracotomy. The heart was excised and retrogradely perfused *via* aortic cannulation inside a temperature-controlled (37 °C) Langendorff perfusion system. A modified Tyrode’s solution (in mM, NaCl 140, KCl 4.7, MgCl_2_ 1.05, CaCl_2_ 1.3, Glucose 11.1, HEPES 10, pH 7.4 at 37 °C) bubbled with 100% O2 was used, and the perfusion pressure was maintained at 70~90 mmHg by adjusting the flow rate between 1 and 2.5 ml/min.

### Pacing strength-duration curve characterization

A platinum bipolar electrode (A-M Systems, catalog number 778000) was used for *ex vivo* studies; and various (1 mm bipolar; 1-, 1.5-2 mm unipolar) graphene electrodes were used for ex- and *in vivo* studies. Electrodes were placed at the anterior center of the left ventricular surface. Stimulating monophasic square pulses with different pulse duration (ms) and pulse amplitude (volt) were generated by a pulse generator (ADInstruments, PowerLab 26T). For mouse (*ex vivo)* and rat (*in vivo)* hearts, the stimulating pacing cycle length of 150 ms (*i.e.*, 400 beats per minute heart rate) and 125 ms (*i.e.*, 480 beats per minute heart rate) were used, respectively. The stimulating pulse amplitude ranged from 0 to 20 V as the pulse generator could afford and was increased in a 0.1 V step. The pacing threshold was defined as the minimal voltage that could pace the heart for at least 10 consecutive beats.

### *Ex vivo* far-field pseudo-electrocardiogram (ECG) and graphene-electrogram (gEG) recording

Both *ex vivo* ECG and gEG signals were acquired using PowerLab 26T and LabChart software (ADInstruments) when the heart was perfused inside the Langendorff chamber. ECG was recorded using a 3-sensing electrodes setup: a positive electrode was placed near the right atrium, a ground electrode was placed near the left atrium, and a negative electrode was placed near the apex of the heart. gEG was recorded using a 2-sensing electrodes setup: a positive electrode (*i.e.*, GET-electrode) was placed on the anterior surface of the heart, and a negative electrode was placed near the left atrium.

### *In vivo* ECG and gEG recording

In vivo ECG and gEG signals were acquired using PowerLab 26T and LabChart software (ADInstruments). ECG was recorded using a 3-sensing electrodes setup in the Lead II configuration: a subdermal positive electrode was placed in the left leg, a subdermal ground electrode was placed in the left arm, and a subdermal negative electrode was placed in the right arm. gEG was recorded using a 2-sensing electrodes setup: a positive electrode (*i.e.*, GET-electrode) was placed on the anterior-lateral surface of the heart, and a subdermal negative electrode was placed on the right arm.

### Optical mapping

Optical mapping was performed as previously described^57^. The cardiac contraction was arrested with 10~13 μM blebbistatin (Cayman Chemicals, catalog number 13186), an electromechanical uncoupler, for 15~20 min before loading the voltage-sensitive dye RH237 (1.25 mg/ml dye stock solution in DMSO, Biotium, catalog number 61018) and calcium-sensitive dye Rhod-2 AM (1 mg/ml dye stock solution in DMSO, Thermo Fisher Scientific, catalog number R1244). The heart was subjected to a 15 min equilibration period followed by dye staining to allow the washout of extra dye. The heart was illuminated by an LED at 520 +/- 17 nm wavelength. Voltage fluorescence signals (filtered by a 695 nm long-pass filter) and calcium fluorescence signals (filtered by a 572 +/- 14 nm band-pass filter) were simultaneously acquired by two CMOS cameras with 100 × 100 pixels resolution (ULTIMA-L, SciMedia) at 2 kHz sampling frequency. The field of view of the camera was 15 × 15 mm. The platinum bipolar electrode or the 1 mm unipolar graphene electrode was placed at the center of the anterior surface of the mouse heart. Electrical stimuli (with 2 ms pulse duration) were applied to determine the pacing voltage threshold. And the heart was paced at 1.5X pacing voltage threshold and 2 ms pulse duration. The pacing cycle length (in ms, 200, 175, 150, 125, 100) was varied to determine the cardiac restitution properties.

### Mouse model for optogenetics

Mice with channelrhodopsin 2 (ChR2) expression in cardiomyocytes alone were created by cross-breeding one parent (Stock No. 011038, Jackson Labs) having Cre recombinase expression driven by the cardiac-specific alpha myosin-heavy chain promoter (αMyHC)^58^ with another flox-ChR2-YFP parent (Stock No. 024109, Jackson Labs)^59^. Genotyping of tail snips (Transnetyx) confirmed the genotype of properly expressing offspring (i.e., αMyHC-Cre-ChR2 mice) to be used in experiments.

### Optogenetic pacing

A blue (470 nm) LED was placed above the heart illuminating the anterior side of the heart. Cardiomyocytes were activated by photons passing through the transparent graphene electrodes. Optical pacing cycle Length (in ms, 200, 150) was faster than the intrinsic heart rate. Pacing and capture were verified in both reference ECG and graphene electrode recordings. Signals were acquired with PowerLab 26T and LabChart software (ADInstruments).

### Optical data analysis

Optical data of transmembrane potential and calcium were analyzed using a custom MATLAB (R2021a) software. Four different parameters were measured. These include (1) action potential duration 80 (APD80) - the time interval from the activation time (*i.e.*, time of maximum 1^st^ derivative of the upstroke) to 80% repolarization, (2) calcium decay time constant (Ca Tau, τ) - a value calculated by fitting an exponential curve 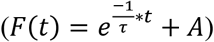 to the calcium transient decay phase (from 30% to 90% decay), (3) transverse and (4) longitudinal conduction velocity (CVT and CVL) were in parallel and perpendicular to the cardiac fiber direction. These two parameters were calculated using differences in activation times and the known interpixel distances.

### *In vivo* rat study

Surgery procedures were performed as previously described^60^. All the surgical procedures were performed under general anesthesia with 3% (v/v) isoflurane inhalation until the animal no longer responded to a toe pinch. The rat was intubated by a 16-gauge cannula and was ventilated using a VentElite small-animal ventilator (Harvard Apparatus, catalog number 55-7040) at a ventilation rate of 80 breaths per minute under pressure control mode with the peak inspiratory pressure limit of 14 cmH2O. After the left thoracotomy, the lung was gently retracted by a cotton swab to expose the heart. The pericardium was gently removed using another cotton swab.

*In vivo* ECG monitoring was performed as aforementioned. For the pacing purpose, a 1 mm bipolar graphene electrode was placed at the anterior-lateral side of the left ventricular surface. Atrioventricular node block (AV block), one kind of cardiac arrhythmia, was induced using a previously described method^57^. Briefly, the rat received (intraperitoneally, IP) 120 mg/kg caffeine (Millipore Sigma, catalog number C0750) and 60 mg/kg dobutamine (Cayman, catalog number 15582) sequentially. Fast pacing (cycle length 100 ms) at the left ventricle was applied for about 15 minutes after drug treatment until AV block was seen. Baseline sinus rhythm was recorded before drug treatment. AV block recording and cardiac rhythm conversion *via* graphene electrode pacing were performed. The PowerLab 26T was used as the energy source to stimulate the heart. After finishing the study, a very deep level of anesthesia was confirmed, the heart was excised, and euthanasia was achieved by exsanguination.

### Statistical analysis

Results are reported as mean +/- standard deviation (SD) unless otherwise noted. Statistical analyses were performed using statistical software (Graphpad Prism, ver 8.4.3). * P < 0.05, ** P < 0.01, *** P < 0.001. For signal-to-noise ratio analysis, intragroup (*i.e.*, ECG vs. gEG) significance was calculated with a nonparametric Wilcoxon test at a significance level of p < 0.05; and intergroup (*i.e.*, 1- vs. 1.5- vs. 2 mm gEG) significance was calculated with a nonparametric Kruskal-Wallis test in conjunction with Dunn’s multiple comparison test at a significance level of p < 0.05. Nonlinear regression analysis was performed on APD80, Ca Tau, CVT, and CVL using the least-squares regression fitting method. An exponential curve *Y* = *Y_M_* – (*Y_M_* – *E*_0_) * *e^-K*X^* was used. Goodness-of-fit was determined by r-squared value > 0.5. The null hypothesis, whether one curve adequately fitted all data sets, was tested with the extra sum-of-squares F test at a significance level of p < 0.05.

## Acknowledgments

This research was supported by NIH grants R21-HL152324, 3OT2OD023848, and R01 HL141470; and Leducq Foundation grant RHYTHM. D.A acknowledges the support of the Office of Naval Research (ONR) under grant number N00014-18-1-2706, and the Temple Foundation Endowed Professorship.

## Author contributions

Z.L., D.K., and I.E. conceived the overall research goals and objectives. Z.L. and D.K. performed proof-of-concept experiments. D.K. and N.L. designed and fabricated GETs and GET arrays. Z.L., S.N.O., and Z.C. conceived and executed the prototyping of the device design. Z.L. performed soft GET-electrodes fabrication. Z.L. performed *ex vivo* experiments. Z.L. and J.L. performed *in vivo* experiments. N.L. and D.K. performed electrochemical impedance measurements. Z.L. and S.G. performed data analysis. D.K. and N.L. performed electrochemical data analysis. Z.L., D.K., N.L., D.A., and I.E. wrote the manuscript. I.E. and D.A. provided funding. All authors reviewed the manuscript before submission.

## Data availability

The data supporting the findings of this study are available within the paper and its supplementary source data files. Data is also available upon request to the corresponding author.

## Code availability

Optical mapping data were processed using a custom MATLAB (R2021a) software (called “OMZer”) that is available at https://github.com/optocardiography/OMZer.

