## Supplementary Information for "Graphene Biointerface for Cardiac Arrhythmia Diagnosis and Treatment"

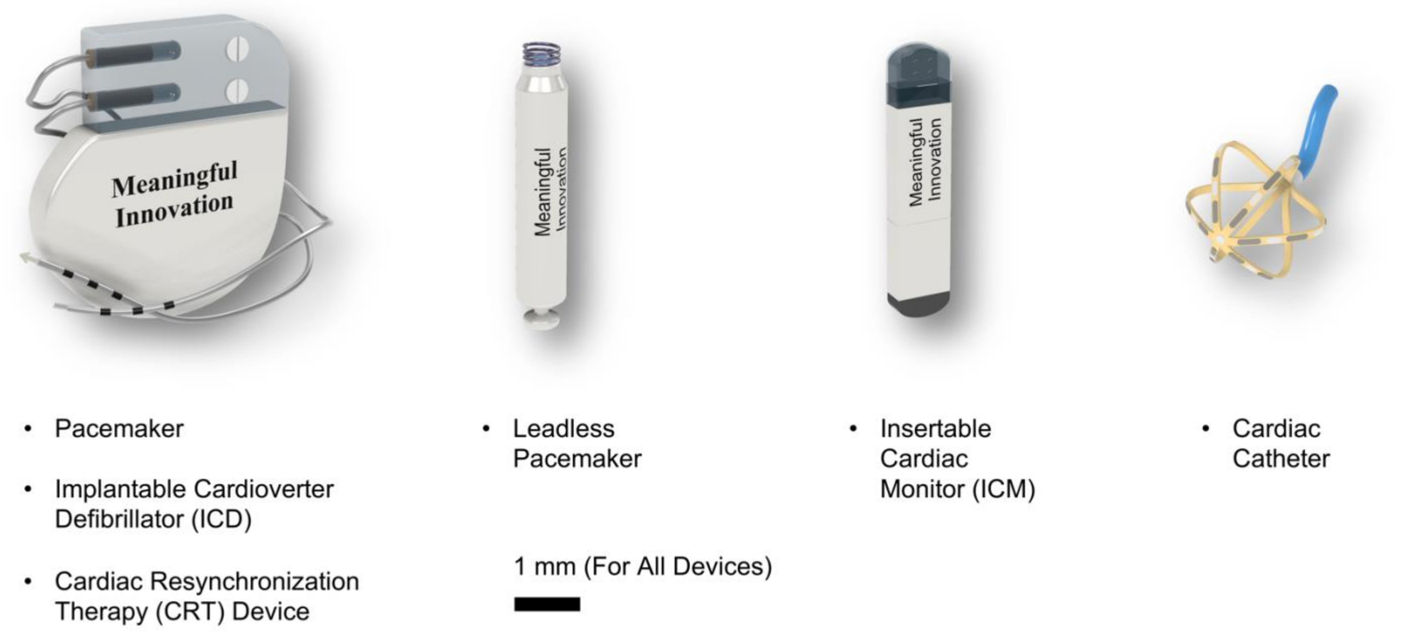


***Figure S1. Examples of currently available clinical devices for cardiac electrophysiology diseases monitor and treatment.***

***Table S1. Examples of clinical devices for cardiac electrophysiology diseases monitor and treatment and potential complications.***

<table>
<thead>
<tr><th>Reference</th><th>Cardiac Electrophysiological Diseases</th><th>Monitor</th><th>Treatment</th><th>Potential Complications[1–10]</th></tr>
</thead>
<tbody>
<tr><td>11–13</td><td>• Atrial Fibrillation<br>• Atrial Flutter</td><td>Insertable Cardiac Monitor (ICM)</td><td>Cardiac Ablation Catheter</td><td rowspan="4">• Infection at The Surgical Site<br>• Device Migration<br>• Erosion of The Device through The Skin<br>• Lead Dislodgement<br>• Lead Failure<br>• Local Pain<br>• Vascular Injury<br>• Haematoma<br>• Cardiac Perforation<br>• Pneumothorax<br>• Deep Vein Thrombosi<br>• Atrio-Esophageal Fistula<br>• Pulmonary Vein Stenosis<br>• Cardiac Tamponade / Hemopericardium<br>• Phrenic Nerve Injury<br>• Stroke</td></tr>
<tr><td>14</td><td>• Sick Sinus Syndrome<br>• Conduction Block</td><td>Implantable Pacemaker</td><td>Implantable Pacemaker</td></tr>
<tr><td>15,16</td><td>• Ventricular Tachycardia<br>• Ventricular Fibrillation</td><td>Implantable Cardioverter Defibrillator (ICD)</td><td>ICD, Cardiac Ablation Catheter</td></tr>
<tr><td>17</td><td>• Heart Failure Followed by Arrhythmia</td><td>Cardiac Resynchronization Therapy (CRT) Device</td><td>Cardiac Resynchronization Therapy (CRT) Device</td></tr>
</tbody>
</table>

### Motivation as to Solve Clinical Issues

Medical devices have been minimizing a lot that benefits the implantation procedures and reduces patient discomfort, however, these cardiac devices and/or electrodes are still in a rigid form which is not ideal to fit the complex mechanical properties of soft tissues. This may constrain natural motion of the heart, injure soft tissues, restrict user mobility, and make patients temporarily or permanently feel discomfort. As a result, research is needed to develop cardiac monitoring and therapeutic devices with improved conformability to soft organs[18–21].

***Table S2. Examples of emergent soft bioelectronic technologies for advanced cardiac electrophysiology.***

| Ref | Function | Transparent | Functional Material | Animal Model (Heart) | Patterning Method | Electrode Number | Device Thickness (µm) |
|---|---|---|---|---|---|---|---|
| This Work | Pacing & Sensing | Yes | Graphene | Mouse (Ex Vivo), Rat (In Vivo) | Biopsy Punch & Mechanical Cutter | 1~4 | 0.5 |
| 22 | | No | Gold-Silver Nanowire | Pig (In Vivo) | Drop Casting | 25 | 400 |
| 23 | Sensing Only | Yes | Silver Nanowire | Mouse (Ex Vivo), Rat (Ex Vivo) | Photolithography | 9 | 25 |
| 24 | | | PEDOT:PSS | Rabbit (Ex Vivo), Pig (In Vivo) | | 64 | 100 |
| 25 | | No | Silicon Nanomembrane | Rabbit (Ex Vivo) | | 396 | 38 |
| 26 | | | Gold | Rabbit (Ex Vivo) | | 34 | 150 |
| 27 | | | Gold | Rabbit (Ex Vivo), Human Donor (Ex Vivo) | | 64 | 180 |
| 28 | | | $Ti_3C_2$ Mxene | Human (ECG, *Wearable*) | $CO_2$ Laser | 1 | 1000 |
| 29 | | Yes | PEDOT:PSS | | Inkjet Printing & Sputtering Deposition | 1 | < 1 |
| 30 | | Yes | Silver Nanowire | | Spin Coating | 2 | 1000 |

Silver (Ag) nanowires have the highest electrical conductivity among metal nanowires[31], but they are susceptible to oxidation and corrosion and the leaching Ag ions may induce adverse health effects[21,32–35].

PEDOT:PSS generally has lower conductivity than metals[36–39], its performance typically drops substantially after immersion in aqueous biological environments[38,39], and the acidity of the PSS chain may cause degradation in devices[40].

MXene is unstable in an open environment even at room temperature and its production cost is still high[41,42].

Gold electrodes are opaque and can exhibit prominent light-induced artifacts during optogenetic stimulation[43].

***Table S3. Examples of reported graphene application for electrophysiology studies compared to this work.***

| Ref | | Function | Animal Model | Patterning Method | Electrode Number | Device Thickness (µm) |
|---|---|---|---|---|---|---|
| This Work | In Vivo, Ex Vivo | Sensing (Electrogram), Pacing (Heart) | Mouse Heart (Ex Vivo), Rat Heart (In Vivo) | Biopsy Punch, Mechanical Cutter | 1~4 | 0.5 |
| 44 | In Vitro | Sensing (Intracellular and Extracellular Action Potential) | Human Embryonic Stem Cell - Derived Cardiomyocytes | Photolithography, Reactive Ion Etching | 80 | --- |
| 45 | In Vitro | Sensing (Extracellular Action Potential) | Embryonic-Rat Cardiac Tissue | Photolithography and Oxygen Plasma Etching | 64 | 13 |
| 46 | In Vivo | Sensing (Electrocorticography), Pacing (Neurons) | Mouse Brain, Rat Brain | Photolithography & Oxygen Plasma Etching | 128 | 3.2 |
| 47 | In Vivo | Sensing (Cortex), Stimulating (Cortex) | Mouse Brain | Photolithography & Oxygen Plasma Etching | 16 | 25 |
| 48 | In Vivo | Sensing (Electrocorticography) | Mouse Brain | Photolithography & Oxygen Plasma Etching | 16 | 58 |
| 49 | In Vivo | Sensing (Electrocorticography) | Mouse Brain | Photolithography & Reactive Ion Etching | 16 | 8 |
| 50 | In Vivo | Sensing (Intracortical Recording) | Rat Brain | Photolithography & Reactive Ion Etching | 64 | 13 |
| 51 | In Vivo | Sensing (Electrocorticography) | Rat Brain | Photolithography & Reactive Ion Etching | 32 | 14 |
| 52 | In Vivo | Sensing (Electrocorticography) | Rat Brain | Photolithography & Reactive Ion Etching | 16 | 12 |
| 53 | In Vivo | Sensing (Electroretinography) | Monkey Cornea, Rabbit Cornea | --- | --- | 7~27 |
| 54 | Wearable | Sensing (ECG) | Human (ECG) | Mechanical Cutter | 1 | < 0.5 |

***Table S4. Comparison table of the state-of-the-art microelectrode arrays and their electrochemical properties.***

| Reference | Functional Material | Electrochemical Parameters | | | Substrate | Thickness | |
|---|---|---|---|---|---|---|---|
| | | Z (1kHz) Ω | CSC mC/cm$^2$ | CIC µC/cm$^2$ | | functional (nm) | total (µm) |
| This work. Large GETs | | 2.5±0.7k | 4.3±0.6 | 83±13 | PMMA | <1 | 0.5 |
| This work. Small GETs | | 5.1±1.6k | 63.7±14.6 | 704±144 | PMMA | <1 | 0.5 |
| 55 | | 286.4±92.6k | 0.0878 | 57.13 | $SiO_2$/Si | <1 | 525 |
| 56 | | 1500k | --- | --- | PET | <1 | 50 |
| 57 | Graphene | 50-600k | --- | --- | Parylene-C | <1 | 15 |
| 58 | | 872.9k | --- | --- | PET | <1 | 50 |
| 59 | | 3000-4000k | --- | --- | $SiO_2$/Si | <1 | 525 |
| 60 | | 170±11.1k | --- | --- | $SiO_2$/Si | <1 | 525 |
| 61 | | 1956±330k | --- | --- | Parylene-C | <1 | 4 |
| 62 | | 908±488k | 0.0224 | --- | Parylene-C | <1 | 4 |
| 63 | Graphene-fiber | 37.9±5.47k | 200±25 | 10340 | --- | 40000 | --- |
| | Graphene | 2650±260k | 0.91±0.13 | 150±50 | | <1 | |
| 64 | Gold/Graphene | 860±70k | 1.58±0.21 | 310±20 | PCB | 100 | 1570 |
| | Au | 1080±460k | 0.73±0.11 | 160±40 | | 100 | |
| [27] | Au | 9.5k | --- | --- | PI | 100-300 | 1-3.3 |
| [23] | AgNW | 53-23k | --- | --- | PET | 120 | 25 |
| 65 | AgNW | 0.06k | --- | --- | none | 100000 | none |
| 66 | rGOx + PtNPs | 180±13k | --- | --- | Glass | --- | 525 |
| 67 | CNT | 11.2±7.6k | 372 ± 56 | 6520 | --- | 46000 | --- |
| 68 | CNTs+PPy | 2.06k | 1244 | 7500 | Pt | ＜50 | 200 |
| 69 | PEDOT+CNTs | 16.2±1.07k | 1.21±0.02 | 1250 | Glass | 352±27 | --- |
| 70 | Pt + modified | 0.188k | 3000 | 24000 | $SiO_2$/Si | 200 | 525 |
| | Pt | 100k | 4.4 | 300 | | | |
| 71 | Pt | 100k | --- | --- | $SiO_2$/Si | 150-200 | 525 |
| 72 | Pt | --- | --- | 54 | $SiO_2$/Si | 0.05 | 525 |
| 73 | Pt nanograss | 15k | --- | 350 | Polyimide | 300 | 10 |
| 74 | Nanoporous Pt + CNT | 2.4k | 1.2 | 3000 | --- | 900 | --- |
| 75 | Pt Microporous | --- | 37.67 | 295.9 | Polyimide | --- | 20 |
| | Pt Sputtered | --- | 3.7 | 81.63 | | | |
| 76 | PEDOT | 0.0233±0.0007k | 75.6 ± 5.4 | 3400 | $SiO_2$/Si | --- | 15 |
| 77 | PEDOT+PtIr | 3k | 125 | 2920 | --- | 75000 | --- |
| [28] | $Ti_3C_2$ Mxene | 0.3692±0.0402k | 306.3±35.2 | 670±30 | PDMS | --- | 1000 |
| 78 | Hydrogel | 500k | --- | --- | $SiO_2$/Si | --- | 525 |

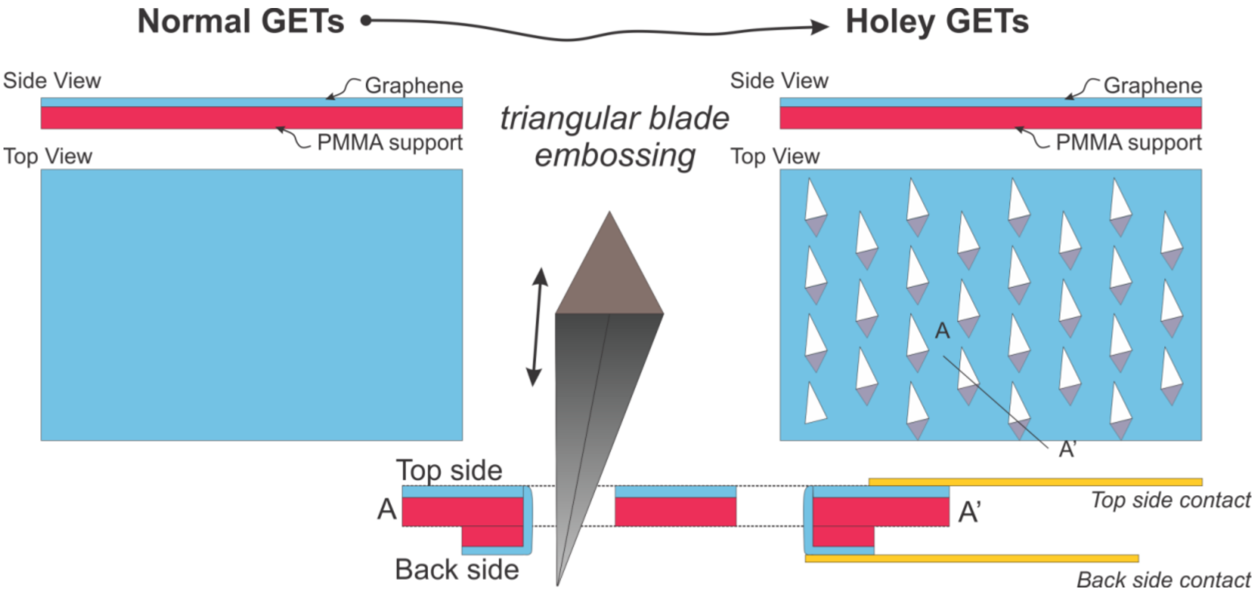


***Figure S2. Schematic of the "Holey GETs" fabrication.*** *Similar to the "classic" GETs, the "holey" ones are made of the graphene/PMMA interface. However, we embossed the surface of the Graphene/PMMA with a Silhouette Cameo cutter. Since this is embossing rather than cutting, the embossed parts of the graphene/PMMA stick to the back-side, exposing larger area of graphene to the back side as well. This opens up need opportunities in terms of building electrical contacts to the GETs not only from the top (graphene) side, but also from the bottom (PMMA) side, The process and electrical characterization of this approach has been published elsewhere*[79,80].

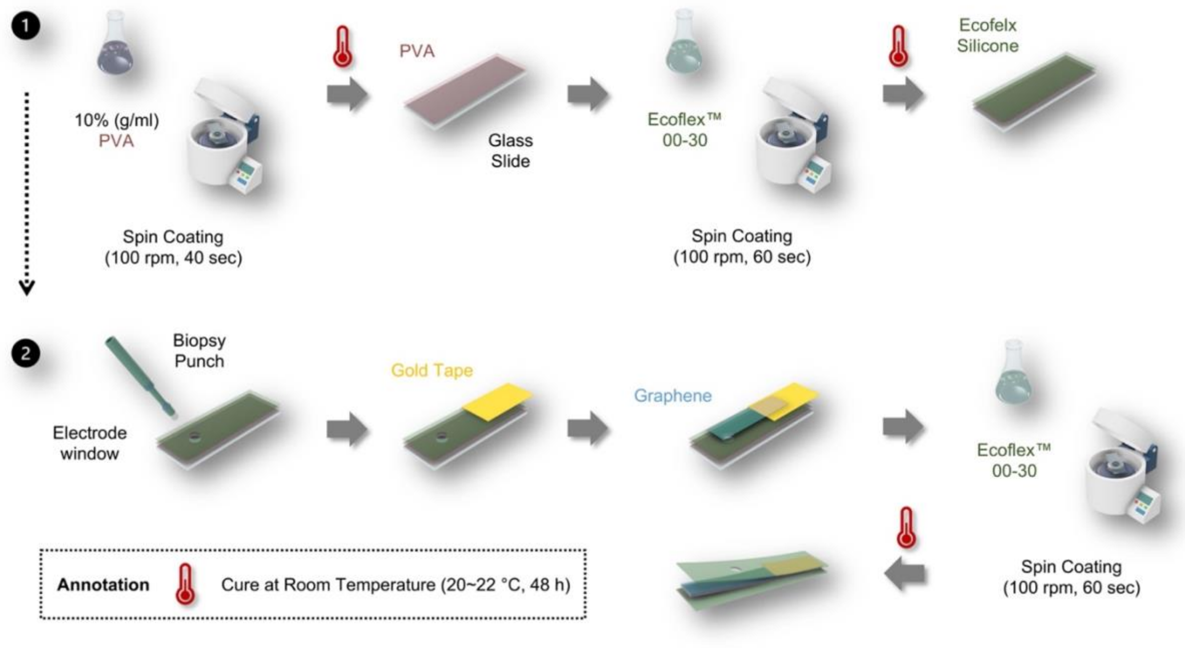


***Figure S3. Schematic of graphene electrodes fabrication.*** *On top of a glass slide, poly(vinyl alcohol) (PVA) (10%, g/ml) is spin coated (100 rpm, 40 sec, cure at room temperature for 48 h) acting as a temporary supporting layer that can ease the delamination of GET-electrode from the glass slide. Then the Ecoflex silicone is spin coated (100 rpm, 60 sec, cure at room temperature for 48 h) acting as a flexible encapsulating layer. Different electrode window patterns are defined on the PVA and silicone layer with biopsy punches of various sizes (diameter 1-3 mm). The gold tape is gently placed onto the encapsulating layer. A graphene tattoo is then placed in such a way that one of its ends covers the electrode window, and the other end contacts the gold tape. Finally, the whole device is encapsulated (spin coated as aforementioned) by one more layer of Ecoflex silicone.*

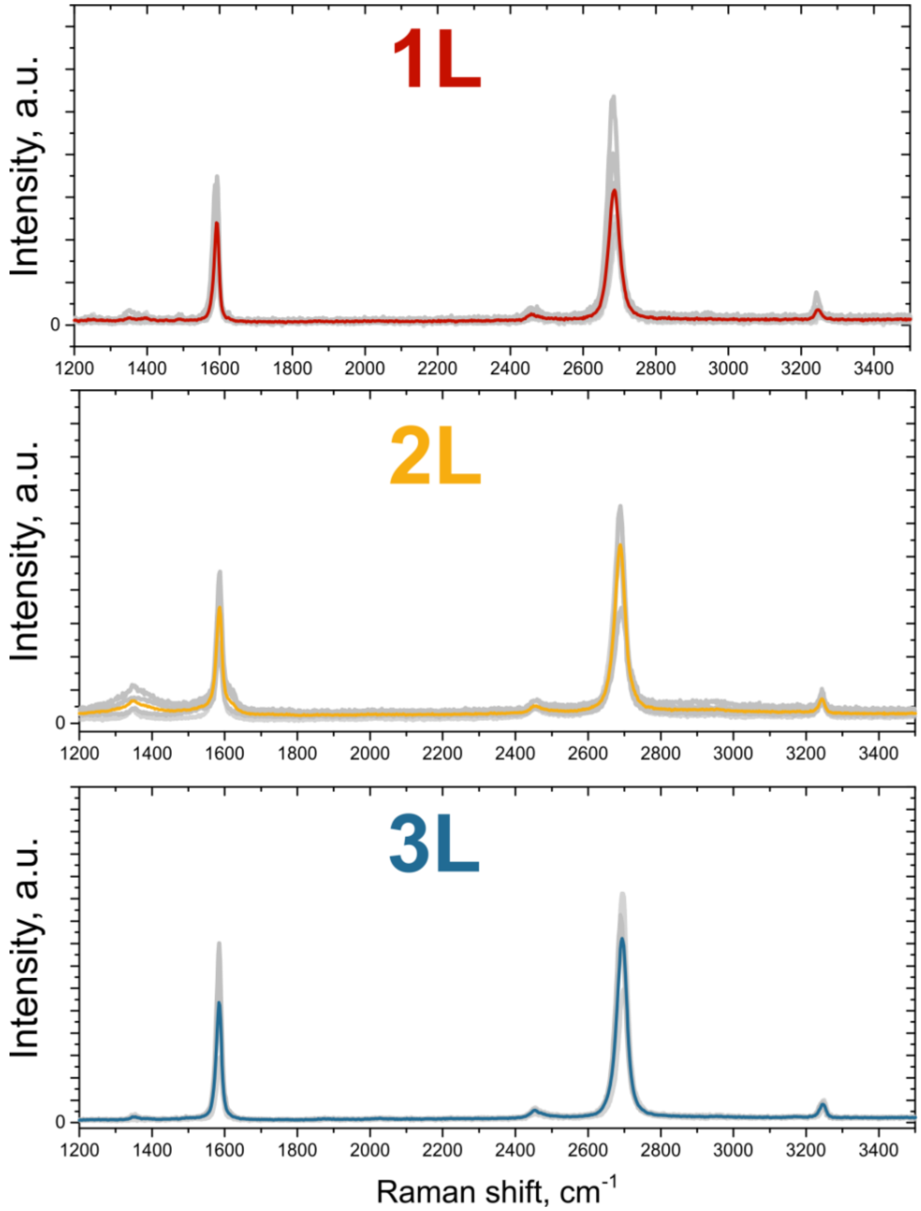


***Figure S4. Raman spectra for the 3 types of GETs reported in this work.*** *In all three cases, the 2D/G peak intensity is in the range of 1.5, indicative of high quality graphene. Gray lines represent individual measurements from N=3 samples, and colored curves represent the average.*

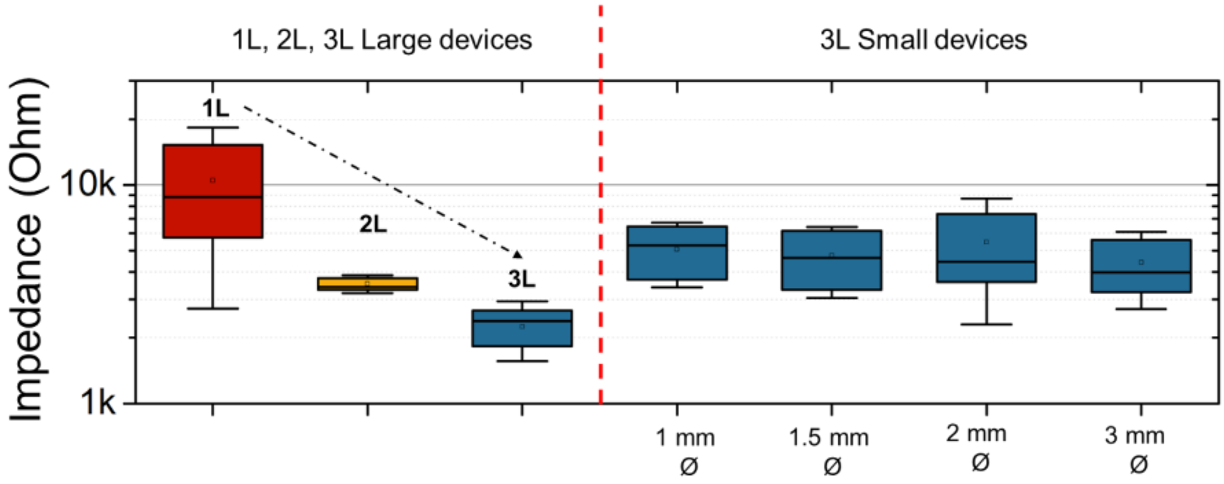


***Figure S5. Box plots of the impedance of large GETs with 1L, 2L, and 3L configurations, and small GETs with 1mm, 1.5mm, 2mm, and 3mm electrode diameters.***

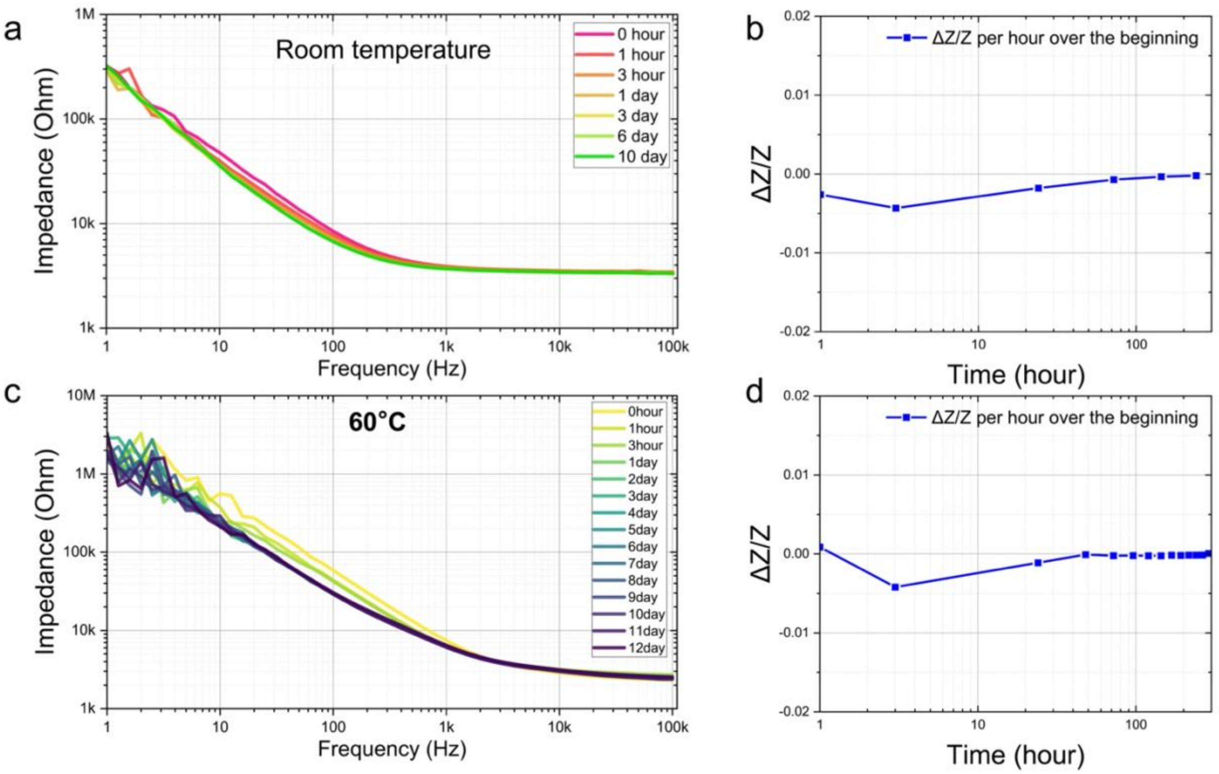


***Figure S6.*** *(a) Bode Impedance plots for GET device measured over a span of 10 days at room temperature kept in 1×PBS, with no visible degradation of the properties. (b) Relatively impedance change per hour over the beginning at 1kHz for the same GET devices (a) at the same measurement. (c) Bode Impedance plots for GET device measured over a span of 12 days at 60℃ kept in 1×PBS, with no visible degradation of the properties. (d) Relatively impedance change per hour over the beginning at 1kHz for the same GET devices (c) at the same measurement. Note: It has been reported that 12 days accelerated aging-test performed at 60 °C is equivalent to 60 days in vivo lifetime. (i.e., with an aging factor of 5 time)*[81].

***Table S5. Quantifiable metrics of the relative change of impedance at 1kHz.***

| | **0 hour** | **1 hour** | **3 hours** | **1 day** | **3 days** | **6 days** | **10 days** |
|---|---|---|---|---|---|---|---|
| **Impedance at 1kHz** | 3884.71 | 3874.59 | 3834.37 | 3718.12 | 3687.06 | 3691.92 | 3700.74 |
| **ΔZ/Z per hour** | - | -0.00261 | -0.00519 | -0.00144 | -0.00017 | 0.00002 | 0.00001 |
| **ΔZ/Z per hour over the beginning** | - | -0.00261 | -0.00432 | -0.00179 | -0.00071 | -0.00034 | -0.00020 |

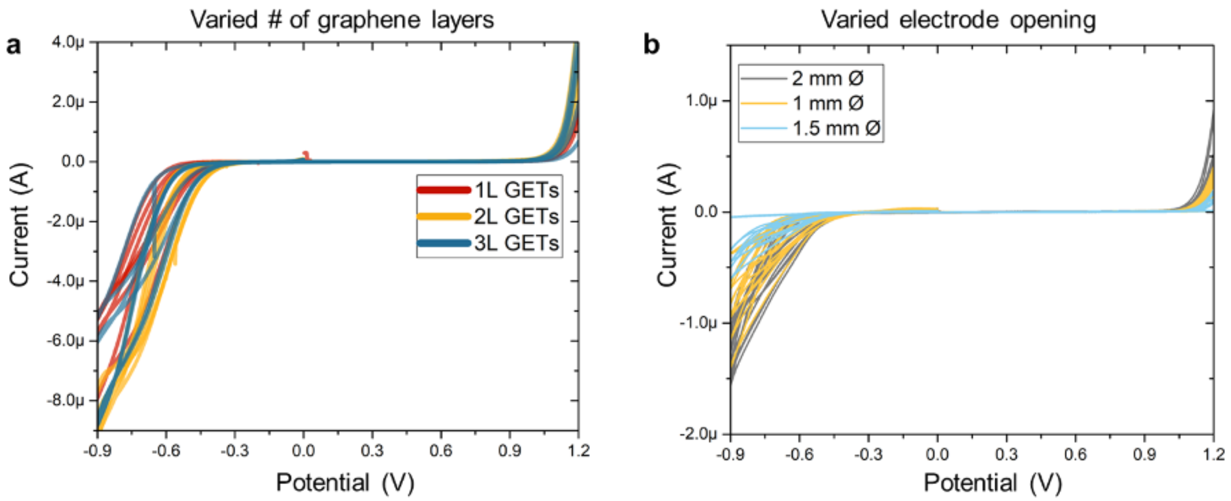


***Figure S7. Cyclic voltammetry of large devices (left) of different # of graphene layers, and small devices (right) of different electrode diameter.*** *No significant difference is found for the 'water window' of the devices regardless of the number of graphene layers and electrode area. Judging from protocol references[81], 10-100 uA of the current could be considered destructive. Although, visually, in the -0.6V to -0.9V range, one may see a raise of the current to ~1-8 uA, it is still way below the 10-100uA, hence still considered safe.*

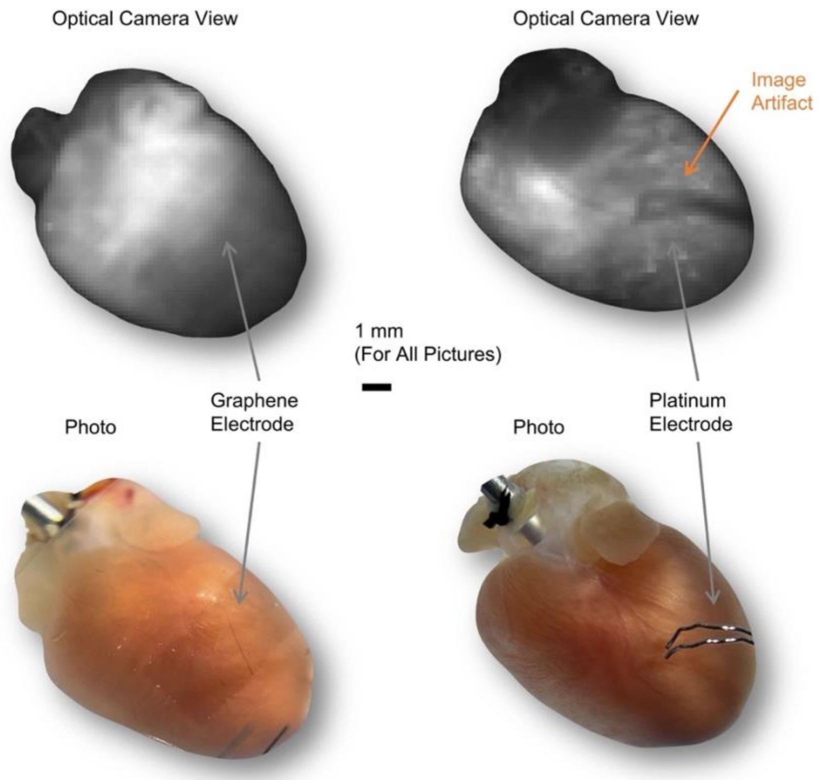


***Figure S8. Comparison views of pacing the heart using a transparent graphene electrode versus using an opaque platinum electrode.*** *A clear image artifact of the platinum electrode can be seen under the optical mapping camera.*

***Table S6. Statistical data on the EIS of the different GETs with different dimensions and # of graphene layers.***

| Device | # of graphene layers | Area | Mean Z ± SD (@1kHz), kOhm | Area-Normalized Mean Z ± SD (@1kHz), Ohm×cm$^2$ |
|---|---|---|---|---|
| Large | 1 | 13-18 mm$^2$ | 10.5±7.8 | 1643±1305 |
| | 2 | 15-18 mm$^2$ | 3.5±0.3 | 609±82 |
| | 3 | 12-18 mm$^2$ | 2.5±0.7 | 345±151 |
| Small | 3 | 3 mm Ø | 4.4±1.7 | 311±120 |
| | 3 | 2 mm Ø | 5.5±3.2 | 172±100 |
| | 3 | 1.5 mm Ø | 4.7±1.7 | 84±30 |
| | 3 | 1 mm Ø | 5.1±1.6 | 40±13 |

***Table S7. Statistical data on the area-normalized CSC.***

| Device | # of graphene layers | Area | Area-Normalized Mean CSC ± SD, $mC/mm^2$ |
|---|---|---|---|
| **Large** | 1 | 13-18 $mm^2$ | 2.8±0.5 |
| | 2 | 15-18 $mm^2$ | 4.3±0.6 |
| | 3 | 12-18 $mm^2$ | 3.7±0.6 |
| **Small** | 3 | 3 mm Ø | 8.1±2.0 |
| | 3 | 2 mm Ø | 15.9±3.6 |
| | 3 | 1.5 mm Ø | 26.5±5.6 |
| | 3 | 1 mm Ø | 63.7±14.6 |

***Table S8. Statistical data on the area-normalized CIC.***

| Device | # of graphene layers | Area | Area-Normalized Mean CIC ± SD, $\mu C/cm^2$ |
|---|---|---|---|
| **Large** | 1 | 13-18 $mm^2$ | 67±26 |
| | 2 | 15-18 $mm^2$ | 83±13 |
| | 3 | 12-18 $mm^2$ | 73±7 |
| **Small** | 3 | 3 mm Ø | 124±29 |
| | 3 | 2 mm Ø | 215±51 |
| | 3 | 1.5 mm Ø | 391±136 |
| | 3 | 1 mm Ø | 704±144 |

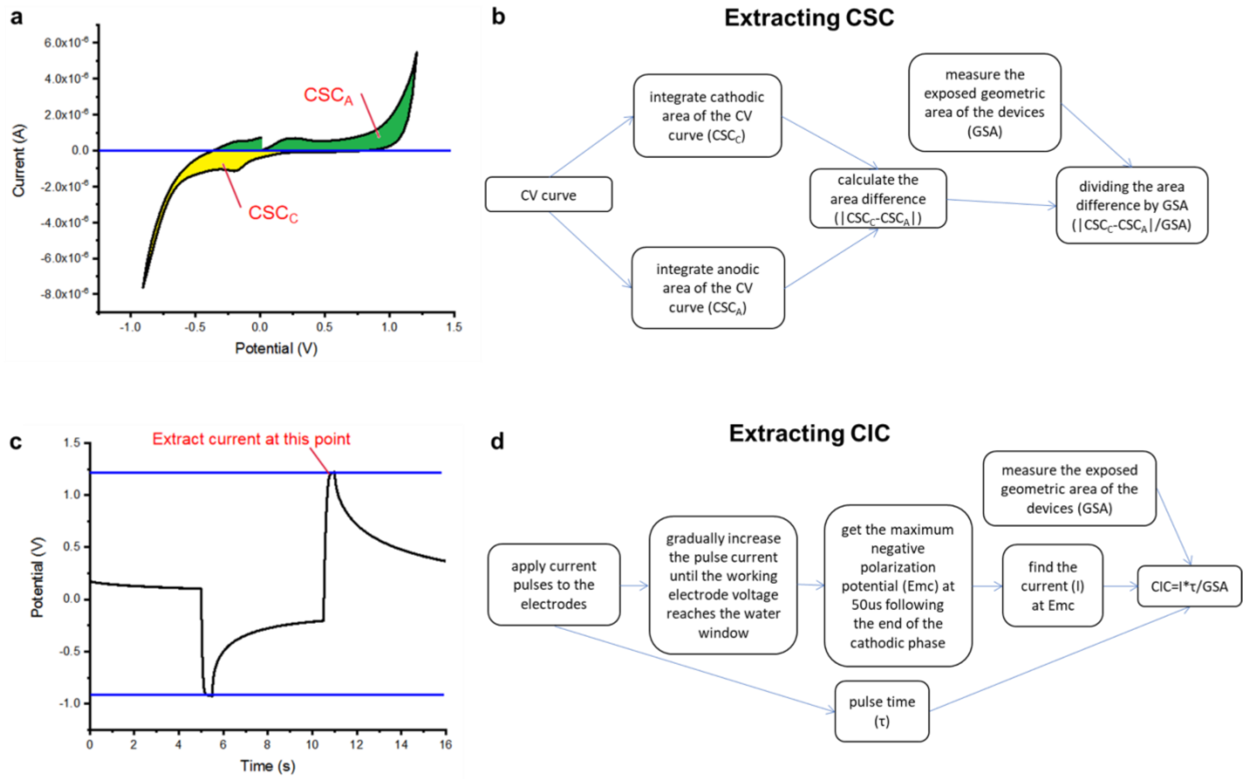


***Figure S9. Schematic and block diagrams of the CSC (a-b) and CIC calculation (c-d).***

*Table S9. Statistical data on the pacing-duration curves.*

| **Pulse Duration (ms)** | **Voltage Threshold**, V (Mean ± SD from 6 hearts for each device kind) | | | | | |
|---|---|---|---|---|---|---|
| | **Mouse Hearts *(Ex vivo)*** | | | | | **Rat Hearts *In vivo*** |
| | **Pt** (bipolar) | 3L-GET, **1 mm Ø** (bipolar) | 3L-GET, **1 mm Ø** (unipolar) | 3L-GET, **1.5 mm Ø** (unipolar) | 3L-GET, **2 mm Ø** (unipolar) | 3L-GET, **1 mm Ø** (bipolar) |
| **10** | 2.03±0.64 | 2.7±1.2 | 1.55±0.5 | 2.5±0.9 | 4.2±0.9 | --- |
| **8** | 2.05±0.64 | 2.9±1.2 | 1.60±0.5 | 2.7±0.7 | 4.6±0.4 | --- |
| **6** | 2.08±0.62 | 3.0±1.3 | 1.7±0.5 | 3.2±0.9 | 4.7±0.6 | --- |
| **4** | 2.10±0.63 | 3.1±1.3 | 1.9±0.7 | 3.7±0.8 | 4.9±0.6 | --- |
| **2** | 2.25±0.69 | 3.5±1.0 | 2.3±0.7 | 4.1±0.9 | 5.4±0.6 | 2.4±1.2 |
| **1** | 2.60±0.76 | 4.4±0.3 | 3.0±0.5 | 4.9±0.4 | 6.1±0.6 | 3.6±0.8 |
| **0.9** | 2.70±0.81 | 4.6±0.4 | 3.1±0.8 | 5.1±0.4 | 6.2±0.6 | 4.4±0.8 |
| **0.8** | 2.80±0.87 | 4.8±0.4 | 3.2±0.9 | 5.2±0.4 | 6.4±0.6 | 4.8±0.8 |
| **0.7** | 2.93±0.87 | 4.9±0.4 | 3.7±0.5 | 5.5±0.4 | 6.6±0.6 | 5.3±0.8 |
| **0.6** | 3.08±0.90 | 5.2±0.5 | 3.9±0.5 | 5.8±0.5 | 6.9±0.6 | 6.5±0.8 |
| **0.5** | 3.27±0.81 | 5.5±0.5 | 4.1±0.4 | 6.2±0.6 | 7.3±0.5 | 8.9±0.4 |
| **0.4** | 3.55±0.88 | 5.9±0.7 | 4.4±0.5 | 6.8±0.7 | 8.0±0.6 | --- |
| **0.3** | 3.9±1.0 | 6.6±0.8 | 5.0±0.5 | 7.9±1.2 | 9.2±0.7 | --- |
| **0.2** | 4.7±1.0 | 8.0±1.0 | 6.3±0.4 | 10.2±3.0 | --- | --- |
| **0.1** | 6.8±1.8 | --- | 9.7±0.6 | --- | --- | --- |

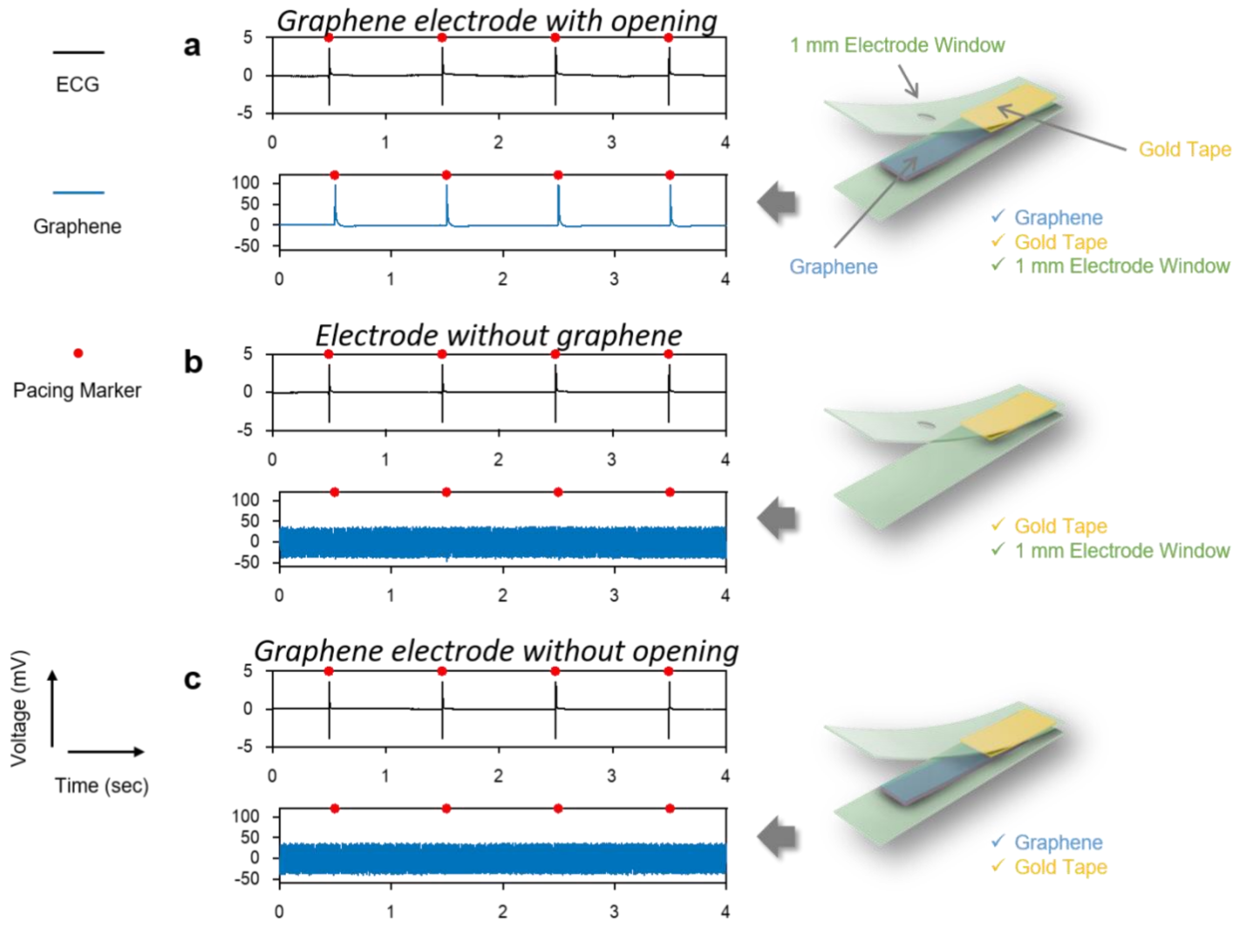


***Figure S10. Representative signals recorded by different graphene electrodes showing good insulation. a**, Simultaneous signal recording using Ag / AgCl electrodes (ECG) and graphene electrodes (graphene). The graphene was well insulated using silicone elastomer except where the electrode window was for the sensing purpose. **b**, Without graphene, even the electrode window was kept, gold tape was still well insulated such that no electrical signals could be recorded. **c**, When the electrode window was removed, no signals could be recorded because of good insulation.*

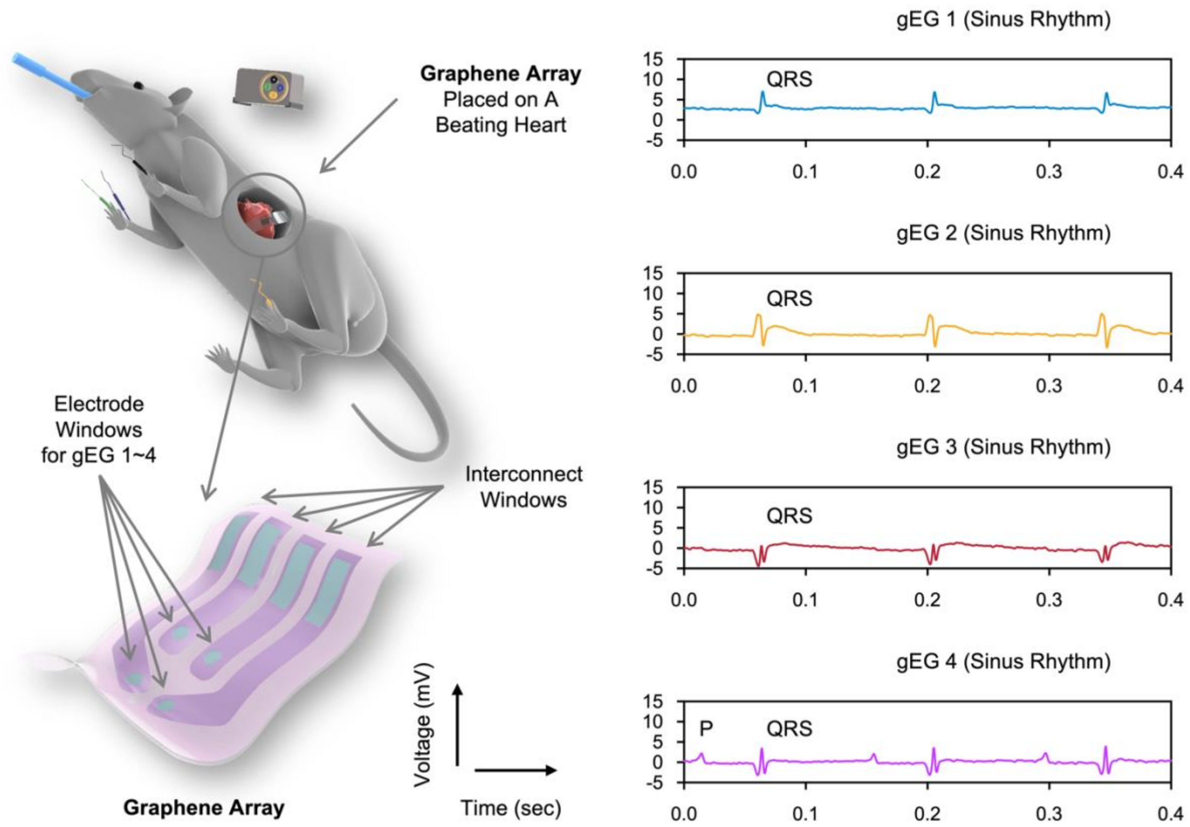


***Figure S11. Cardiac electrogram simultaneously recoded from four different locations of the heart using a graphene array electrode.***
